# Fast synaptic-like axo-axonal transmission from striatal cholinergic interneurons onto dopaminergic fibers

**DOI:** 10.1101/2022.03.25.485828

**Authors:** Paul F. Kramer, Samuel Brill-Weil, Alex C. Cummins, Renshu Zhang, Gisela A. Camacho-Hernandez, Amy H. Newman, Mark A. Eldridge, Bruno B. Averbeck, Zayd M. Khaliq

## Abstract

Transmission from striatal cholinergic interneurons (CINs) controls dopamine release through nicotinic acetylcholine receptors (nAChRs) on dopaminergic axons. Anatomical studies suggest that cholinergic terminals signal predominantly through non-synaptic volume transmission. However, the influence of cholinergic transmission on electrical signaling in axons remains unclear. We examined axo-axonal transmission from CINs onto dopaminergic axons using perforated-patch recordings which revealed rapid spontaneous EPSPs with properties characteristic of fast synapses. Pharmacology showed that axonal EPSPs (axEPSPs) were mediated primarily by high-affinity α6-containing receptors. Remarkably, axEPSPs triggered spontaneous action potentials locally in dopaminergic axons, suggesting these axons perform integration to convert synaptic input into spiking, a function associated with somatodendritic compartments. We investigated cross-species validity of cholinergic axo-axonal transmission by recording dopaminergic axons in macaque putamen and found similar axEPSPs. Thus, we reveal that fast synaptic-like neurotransmission underlies cholinergic signaling onto dopaminergic axons, providing insight into how nicotinic receptors shape electrical signaling directly in axon terminals.

## INTRODUCTION

Interactions between cholinergic and dopaminergic signaling in the striatum have been proposed to play roles in reward, motivation and motor learning (Aosaki et al., 1994; Collins et al., 2016; Ding et al., 2010; Howe et al., 2019), and their imbalance is thought to contribute to the symptoms of Parkinson’s Disease (Lim et al., 2014; Tanimura et al., 2019; Tanimura et al., 2018). Previous work has shown that dopamine release can be modulated by striatal cholinergic interneurons (CINs) through the activation of nicotinic acetylcholine receptors (nAChRs) expressed on the axon shaft and at axon terminals of midbrain dopamine (DA) neurons (Sulzer et al., 2016; Zhou et al., 2002). Studies have shown that activation of CINs using optogenetics (Cachope et al., 2012; Threlfell et al., 2012), stimulation of cortical inputs (Adrover et al., 2020; Kosillo et al., 2016) or spontaneous CIN activity (Yorgason et al., 2017; Zhou et al., 2001) can trigger release from DA axons through a mechanism that occurs independently of somatic action potential generation. Similarly, behavioral studies have suggested that DA signaling in striatal projection axons can occur independently of somatic firing (Mohebi et al., 2019) and have proposed that axonal nAChRs may underlie this function (Mohebi and Berke, 2020). These observations demonstrate that nAChRs on DA axons provide a potentially powerful means of controlling striatal DA release and may enable input-output signaling in the axon terminals that occurs separately from somatic activity. However, little is known of the precise synaptic or non- synaptic functional mechanisms enabling these processes.

While nAChR modulation of DA release has been explored extensively, most studies have focused on CIN-evoked DA release which has left a substantial gap in our understanding of how cholinergic transmission directly influences DA axons. CIN-evoked DA release involves a disynaptic signaling cascade that integrates multiple processes into a single DA measurement. Factors that contribute to this process include calcium-dependent release for both acetylcholine (ACh) and DA, neurotransmitter diffusion and regulation by transporters and degradation enzymes, activation, and desensitization of nAChRs, as well as the voltage-gated ion channels that regulate axonal excitability. It is therefore challenging to infer the specific independent functions of any one factor, such as the dynamics of nAChR activation on DA axons following physiological release of ACh.

Although direct functional data for cholinergic transmission onto DA axons are lacking, predictions regarding ACh-DA interactions have been based on structure data from electron microscopy. Axonal nAChRs on dopaminergic fibers have only rarely been observed in close apposition to presynaptic release sites (Aznavour et al., 2003; Contant et al., 1996; Jones et al., 2001). An earlier immunocytochemical EM study examining identified cholinergic and DA axons did not identify classic synaptic specializations but found numerous close contacts between dopaminergic and cholinergic varicosities (Chang, 1988) suggesting potential functional interactions. Therefore, structural studies find that CIN terminals lack morphological characteristics that narrowly define a synapse and thus suggest that transmission likely occurs through slow, volume-based diffusion. However, how closely the kinetics and frequency dependence of cholinergic transmission onto DA axons is constrained by the known architecture of ACh and DA positive contacts is unclear.

Here, we examine cholinergic transmission onto axonal nAChRs using direct recordings from striatal DA axons of mice and primates in brain slices that separate the axon from the soma and dendrites. We found that CIN-mediated activation of nAChRs resulted in fast, phasic depolarizations in the axons of dopamine neurons that closely resembled nAChR-mediated excitatory post-synaptic potentials (EPSPs) recorded in somas and dendrites of cells in cortex and thalamus (Bennett et al., 2012; Obermayer et al., 2019; Sun et al., 2013). We demonstrate that axonal EPSPs (axEPSPs) are regulated by acetylcholinesterase (AChE), involve α6-containing nAChRs, and exhibit short-term depression that does not involve nAChR desensitization. Interestingly, we show that spontaneous axEPSPs continue to occur even in the absence of spike-mediated release, a characteristic that is shared with point-to-point synapses. Finally, we show that in a subset of axonal recordings, summation of nicotinic axEPSPs triggered spontaneous action potentials (APs) that were initiated locally in the DA axon terminal arborizations. In sum, our direct axonal recordings show that nicotinic receptors control electrical signaling in dopaminergic axons through a mechanism involving fast local activation of α6 nicotinic receptors, providing new insight into the critical role of nicotinic receptors in controlling striatal dopamine release.

## RESULTS

### Spontaneous EPSP-like nicotinic depolarizations recorded in DA axons

Subthreshold voltage dynamics were measured in DA axons located within the dorsomedial striatum. DA axons were identified using tissue slices prepared from dopamine transporter (DAT)-Cre mice in which the red fluorescent protein tdTomato was used to selectively label dopaminergic neurons (**Figure 1A**). To measure the axonal membrane potential, we performed perforated-patch recordings from the cut ends of axons (blebs) located on the surface of the slice (Shu et al., 2007). Perforated-patch recording was used to prevent the rapid rundown of nicotinic currents as reported in cell bodies of dopamine neurons (Wu et al., 2004). In some experiments, the patch was ruptured to allow filling of the axonal cytoplasm with neurobiotin. Post-hoc visualization confirmed that the recorded axons were indeed terminal arbors that had been isolated from the main axon trunk during the slice procedure (**Figure 1B**).

**Figure 1.**
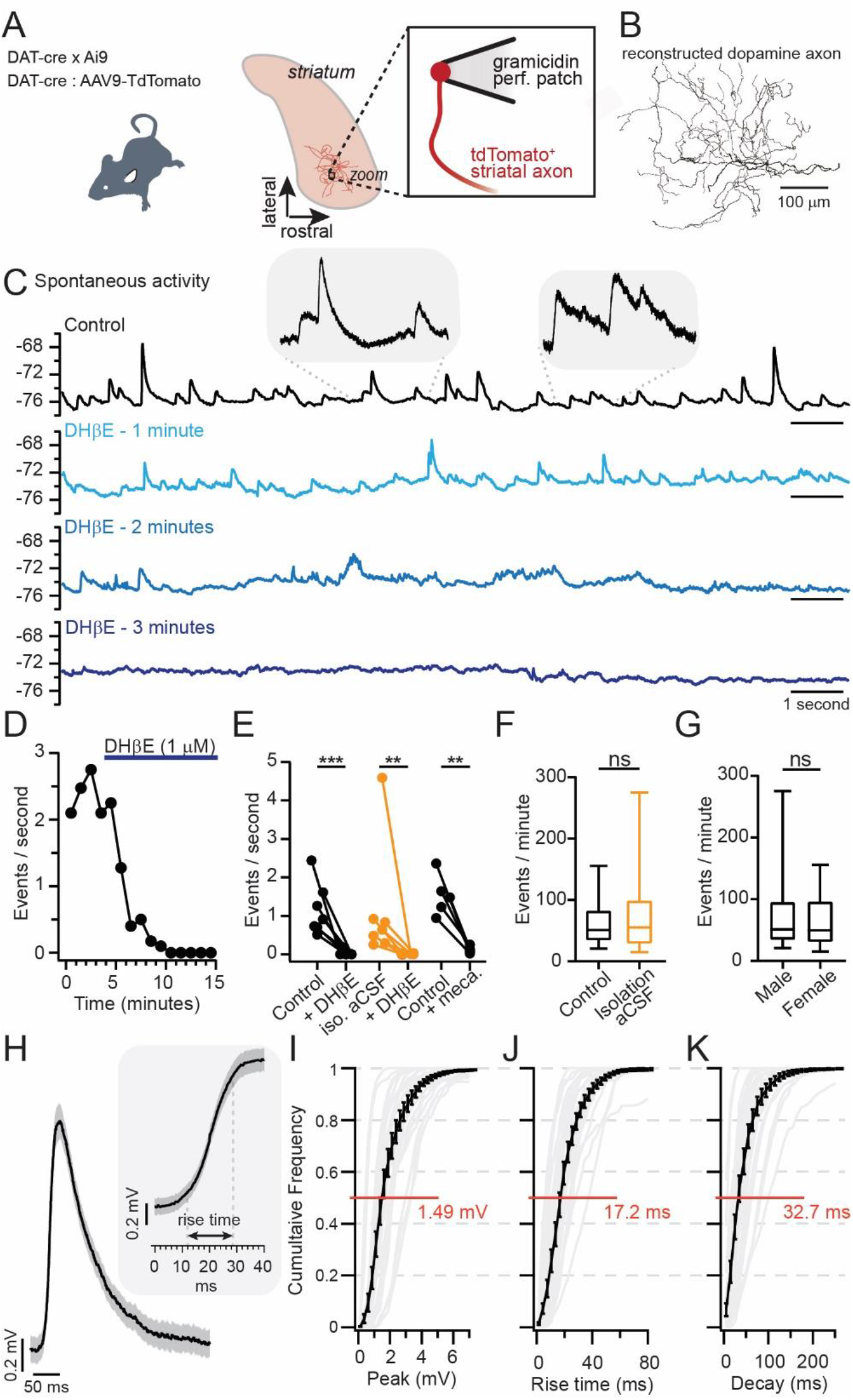
Spontaneous nicotinic axEPSPs recorded in DA axons. **A.** DAT-cre/Ai9 mice and DAT- cre mice infected with AAV9-TdTomato into the SNc were used to visualize DA axons. **B.** Morphological reconstruction of a DA axon filled with neurobiotin. **C.** Spontaneous axEPSPs recorded in perforated patch configuration from a dopaminergic axon in the dorsomedial striatum, control (top) and during DHβE application. **D.** Binned time course of the frequency of axEPSPs shown in (**C**) **E.** Compiled axEPSP frequencies in control (average of baseline before drug wash-in) and drug (average after 8 minutes of wash-in). Black traces recorded in control aCSF, and orange traces recorded in isolation aCSF. Mecamylamine (10 µM) abbreviated as “meca”. **F.** Comparison of spontaneous axEPSP frequency in control aCSF versus isolation aCSF. **G.** Comparison of spontaneous axEPSP frequency in male versus female mice. **H.** Average spontaneous axEPSP recorded from one dopaminergic axon (average of 334 events, error bars ± s.e.m.). Cumulative histograms for the distribution of axEPSP peak amplitudes (**I.**, n=28), rise times (**J.**, n=25), and decay (**K.**, n=27). ** (p < 0.001), *** (p < 0.0001), ns (p > 0.05)

Perforated-patch axonal recordings revealed the presence of spontaneous excitatory postsynaptic potential (EPSP)-like depolarizations in the membrane potential of DA axons. As shown in the **Figure 1C** example, the axEPSPs were small amplitude, phasic events that occurred at a low frequency of 2-3 Hz. AxEPSPs occurred at a median of 51.2 events per minute (n=27), had a median amplitude of 1.49 mV, a median rise time of 17.2 ms, and a median decay time of 32.7 ms (n= 27; **Figure 1H-K**). By contrast, axEPSPs were never observed in recordings from the main axon trunk that is connected to the cell body (Kramer et al., 2020), supporting the idea that axEPSPs are generated in the axon terminals.

AxEPSPs were completely abolished by dihydro-β-erythroidine hydrobromide (DHβE, 1 µM), a nicotinic receptor antagonist with modest selectivity for receptors containing the α4β2 subunit (Gopalakrishnan et al., 1996; Harvey et al., 1996) (**Figure 1C-E**). The axEPSPs were also inhibited by the broad nicotinic receptor antagonist mecamylamine (10 µM; **Figure 1E**) further confirming their nicotinic identity. We saw similar results when we recorded in solutions designed to isolate ACh transmission (isolation aCSF) that contained antagonists against dopamine D2, muscarinic, GABA-B, GABA-A, AMPA and NMDA receptors (**Figure 1E,F**; control: 51.2 events / minute, n = 27; isolation aCSF: 55.4 events/minute, n = 19; Mann-Whitney U = 254, p = 0.965). Furthermore, there was also no difference in the frequency of events observed between male and female mice (**Figure 1G**; Female: 49.8 events / minute, n = 28; Male: 51.2 events/minute, n = 13; Mann-Whitney U = 164, p = 0.628). Therefore, we find spontaneous and phasic nicotinic input mediated by α4β2-containing receptors that appears synaptic in nature.

### Miniature nicotinic axEPSPs recorded in DA axons

Striatal cholinergic interneurons are spontaneously active neurons that are thought to be the sole source of ACh transmission onto DA axons (Brimblecombe et al., 2018). We therefore tested whether the spontaneous axEPSPs are driven by spontaneous cellular activity using tetrodotoxin (TTX) to block sodium dependent APs. Bath application of TTX reduced the frequency of spontaneous axEPSPs (control: 0.81 ± 0.11 Hz; TTX: 0.17 ± 0.05 Hz; **Figure 2A-B**). However, the events were not completely abolished. We observed miniature nicotinic axEPSPs that persisted in the presence of TTX and were blocked by the further addition of DhβE (**Figure 2A-B**). The rise time of axEPSPs were similar between TTX and control conditions (rise time in control: 16.8 ± 0.38 ms vs TTX, 16.2 ± 0.83 ms; p = 0.08, n=9; **Figure 2D**), while the amplitude was reduced (control: 1.59 ± 0.02 vs TTX, 1.31 ± 0.04 mV; p<0.0001, n=9; **Figure 2F**) which predominantly affected large amplitude events (**Figure 2G**). Interestingly, the decay kinetics of the miniature events were substantially faster in the presence of TTX, from a time constant of 33.4 ± 0.70 in control to 26.6 ± 0.92 ms in TTX (p<0.0001, n=9; **Figure 2E**). This suggests that TTX-sensitive sodium channels in DA axons likely boost and prolong the decay of axEPSPs. Furthermore, the presence of miniature axEPSPs may be indicative of close apposition between a set of nAChRs and vesicle release sites.

**Figure 2.**
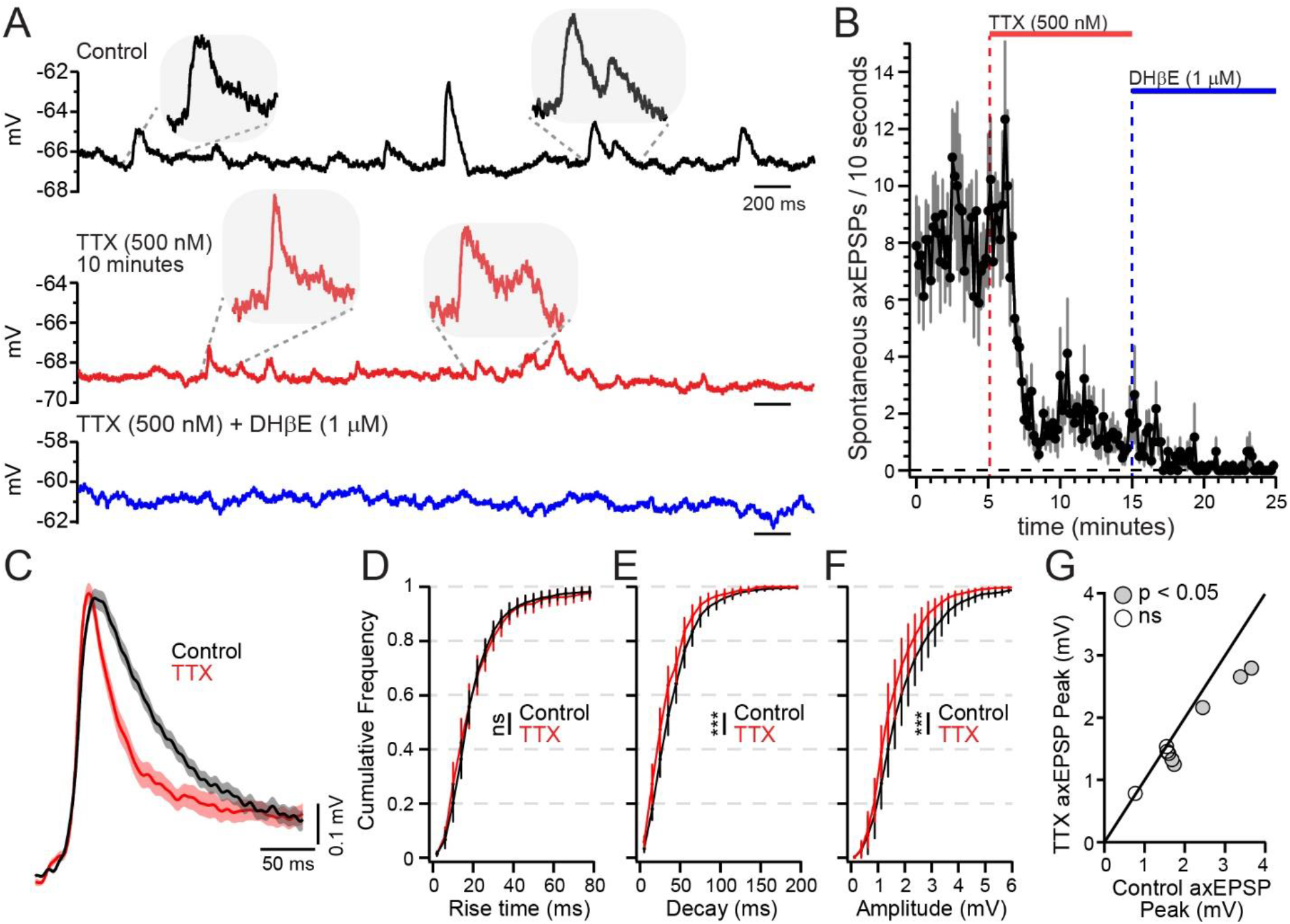
Spontaneous ‘miniature’ nicotinic axEPSPs recorded from DA axons. **A.** Spontaneous axEPSPs measured in control (top), in the presence of tetrodotoxin (TTX, 500 nM, middle), and TTX plus DHβE (1 µM). Example traces are from the same axon. **B.** Time course of spontaneous axEPSP frequency (events per 10 seconds) in control, TTX and DHβE (n = 9 axons). **C.** Event averages from one axonal recording of axEPSPs in control (average of 358 events) and miniature axEPSPs in TTX (average of 176 events; ± s.e.m in shaded region). Cumulative histograms comparing control and TTX axEPSPs in rise time (**D**), decay (**E**), and peak amplitude (**F;** n = 9 axons). **G.** Average axEPSP event amplitudes in control plotted against average in TTX. Each point is the average from all control or TTX event amplitudes plotted against each other from one recording. Solid line represents a line of unity.

### Stimulated nicotinic axEPSPs in DA axons

We next tested evoked release from cholinergic interneurons onto DA axons. To test this, we crossed ChAT-Cre with DAT-Cre mouse lines (ChAT-Cre/DAT-Cre). Using AAVs to drive protein expression in selected cell types, we expressed channelrhodopsin (ChR2) and tdTomato in cholinergic interneurons and midbrain dopamine neurons, respectively (**Figure 3A**). Light stimulation of CINs resulted in optically-evoked phasic depolarizations in the dopamine axons (**Figure 3B**). Optically-evoked axEPSPs, with the LED power tuned just below the threshold of evoking an axonal AP in the DA axon, had a mean amplitude of 8.5 ± 1.66 mV and were inhibited by DHβE (**Figure 3C**). We next measured electrically-stimulated cholinergic transmission using a bipolar electrode placed in the dorsomedial striatum and recorded in cholinergic isolation aCSF (**Figure 3D**). Electrically evoked axEPSPs, also tuned below the threshold of evoking an axonal AP in the DA axons, had a mean amplitude of 5.65 ± 0.76 mV with a coefficient of variation (CV) of 0.16 ± 0.03 (n = 16). The kinetics of electrically evoked axEPSPs were similar to those that were optically evoked (rise time: o-stim = 13.3 ± 2.6 ms; e- stim = 12.1 ± 1.8 ms; U=57, p = 0.90), and were also blocked by DhβE (**Figure 3E-F**). Importantly, the kinetics and latency to onset of the electrically-evoked axEPSPs did not differ in control versus cholinergic isolation solutions, consistent with direct release of ACh onto dopamine axons, and with a lack of presynaptic inhibition of ACh release under these conditions by either dopamine D2 receptor or muscarinic M2 receptors (**Figure 3G-K**; rise time: ctrl = 12.1 ± 1.8 ms, iso = 15.5 ± 1.8 ms, U = 155, p = 0.38; onset: ctrl = 5.93 ± 0.37 ms, iso = 6.13 ± 0.42 ms, U = 146, p = 0.34; decay: ctrl = 71.5 ± 17.2 ms, iso = 123 ± 33.3 ms, U = 141, p = 0.28; amplitude: ctrl = 4.9 ± 0.48 mv, iso = 4.2 ± 0.50 mV, U = 159, p = 0.57).

**Figure 3.**
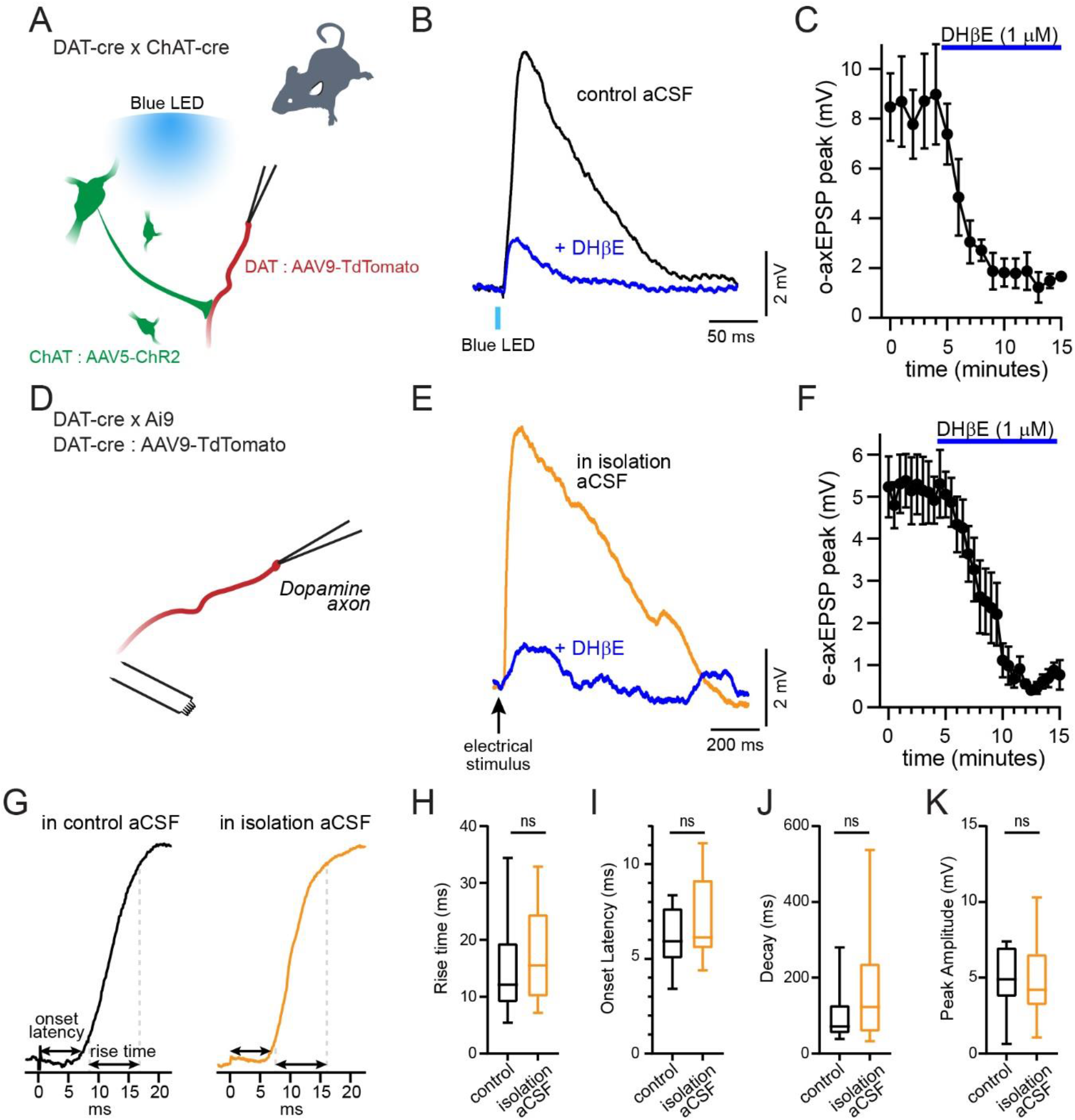
Stimulated axEPSP in DA axons. **A.** Experimental setup. DAT-Cre x ChAT-Cre double transgenic mice were injected with cre-dependent ChR2 (AAV5) into the dorsal medial striatum, and cre-dependent tdTomato into the SNc. **B.** Example traces of optogenetic activation of ChAT+ interneurons while recording from DA axons in control (black) and after DHβE (blue) **C.** Time course showing the inhibition of the optically stimulated axEPSP in DA axons by DHβE (1 µM, n = 6 axons). **D.** Experimental setup. DA axons were identified by expression of tdTomato. A bipolar electrode was used for electrical stimulation. **E.** Example traces of electrically stimulated axEPSPs in isolation aCSF (orange) and following DHβE (blue). **F.** Time course showing the inhibition of the electrically stimulated axEPSP in DA axons by DHβE (1 µM, n = 7 axons). **G.** Zoomed-in portion of stimulated axEPSPs to show onset latency and rise time of example traces in control aCSF (black) and isolation aCSF (orange). Comparison between control (n = 17 axons) and isolation (n = 21 axons) aCSF for axEPSP rise times (**H.**) onset latency (**I.**) decay (**J.**) and peak amplitude (**K.**)

### α6-containing nAChRs account for a majority of the axEPSP

Alpha-6 subunit containing nAChRs are expressed on the axons of dopaminergic neurons (Zoli et al., 2002) where they modestly contribute to nicotinic receptor-dependent dopamine release in the dorsal striatum (Exley et al., 2008). Yet the functional effect of this subunit on subthreshold nicotinic depolarization in the axon is unknown. Whole-cell somatic recordings of nAChR currents in DA neurons have yielded conflicting data, with one study showing evidence for about 20% of the total nicotinic current blocked by the α6-specific receptor antagonist α- conotoxin-MII (αCtx-MII) (Champtiaux et al., 2003), while another study found no inhibition by the same antagonist (Wu *et al*., 2004). We therefore sought to establish the contribution of α6 subunits to nAChR-mediated membrane depolarization in DA axons.

We tested the effect on axonal nicotinic signaling of conotoxin-P1a (Ctx-P1a), an antagonist that has a high selectivity for α6 over the α3 subunit (Dowell et al., 2003; Gotti et al., 2010; McIntosh et al., 2004). Ctx-P1a (1 µM) reduced the amplitude of the electrically-stimulated axEPSP to 43.8 ± 8.48% of the control amplitude (**Figure 4A**; t(7) = 6.63, p = 0.0006). With subsequent application of DhβE, the axEPSP amplitude was further reduced to 2.75 ± 1.26% of the control value (**Figure 4A**; t(7) = 4.79, p = 0.002). Interestingly, the rise time of the axEPSP was significantly slower when measured in the presence of Ctx-P1a relative to control conditions (**Figure 4B,C**; rise time, control: 14.4 ± 3.4 ms; Ctx-P1a, 23.5 ± 2.7 ms). Examining the effects of Ctx-P1a on spontaneous axEPSPs (**Figure 4D-H**), we found that the frequency of spontaneous axEPSPs was reduced to 30.7 ± 12.9% in the presence of Ctx-P1a (**Figure 4H**), which was further reduced by DHβE to 1.91 ± 1.05%. Of note, the remaining events in Ctx-P1a had smaller amplitudes in the upper quartile of events (control: 0.94 ± 0.02 mV; Ctx-P1a: 0.91 ± 0.02 mV; p = 0.0001, n = 8, **Figure 4F**) and longer rise times overall (control: 18.8 ± 0.2 ms; Ctx- P1a: 20.1 ± 0.24 ms; p = 0.0009 n=8, **Figure 4G**), similar to evoked axEPSPs following Ctx-P1a treatment. These results show that α6-containing receptors are necessary for the majority of the large amplitude spontaneous nicotinic axEPSPs in DA axons.

**Figure 4.**
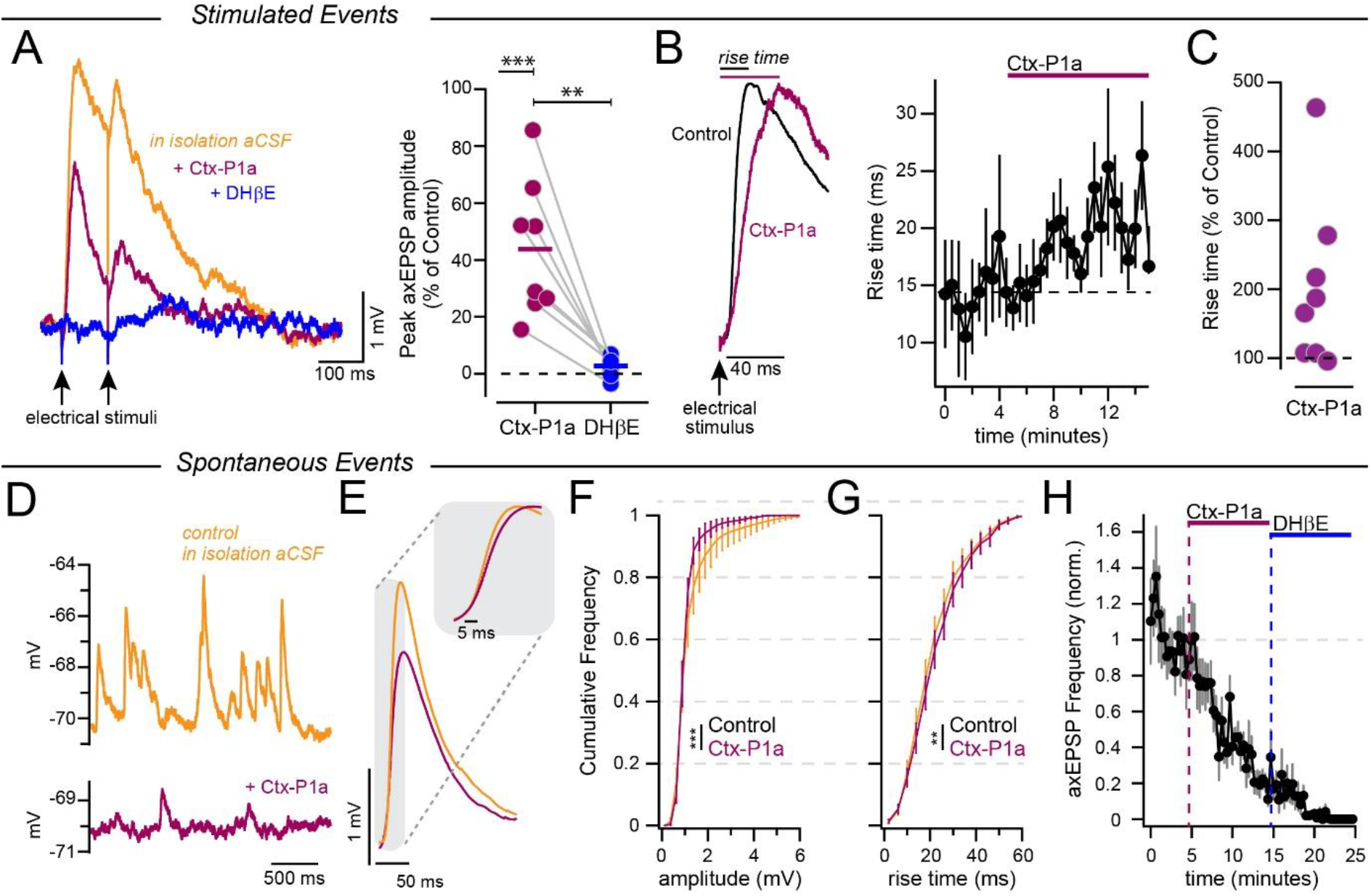
α6-containing nAChRs account for a majority of the axEPSP. **A.** Left panel: Example trace of a stimulated axEPSP in isolation aCSF (orange), conotoxin-P1a (maroon) and DHβE (blue). Right panel: Peak amplitude as a percentage of each axon’s control following conotoxin- P1a (maroon) and DHβE (blue), n = 8 axons. **B.** Left panel: zoomed-in portion of stimulated axEPSPs to show rise time of example traces in isolation aCSF (orange) and with the addition of conotoxin-P1a (maroon). Right panel: Time course of the axEPSP rise time in experiments where conotoxin-P1a was applied (n = 8 axons). **C.** Averaged rise time values for each axon in conotoxin-P1a, plotted as a percentage of the control value. **D.** Example trace of a spontaneous axEPSP in isolation aCSF (orange) and conotoxin-P1a (maroon). **E.** Averaged traces of spontaneous axEPSPs from one axonal recording of control in isolation aCSF (average of 1094 events) and in conotoxin-P1a (average of 700 events, ± s.e.m in shaded region). Cumulative histogram of the event amplitudes (**F.**) and rise times (**G.**) in isolation aCSF (black) and in conotoxin-P1a (maroon; n = 8 axons). **H.** Frequency of spontaneous events (normalized), n = 8 axons.

### Desensitization of nAChRs contributes little to short-term plasticity

Short-term plasticity of neurotransmission is a key factor in the transfer of information between neurons. Previous work examining cholinergic transmission onto muscarinic receptors on striatal medium spiny neurons observed significant depression (Mamaligas et al., 2016) which is consistent with data from voltammetry studies examining CIN-evoked dopamine release (Rice and Cragg, 2004; Shin et al., 2017; Threlfell *et al*., 2012; Zhang and Sulzer, 2004). These data incorporate both dopamine axon and CIN axon release properties, leaving the short-term plasticity of ACh release from striatal CINs onto DA axons unknown.

To directly assess the short-term plasticity of ACh release onto DA axons, we electrically-evoked pairs of axEPSPs at varying inter-stimulus intervals. Responses were measured in isolation aCSF to remove confounding effects of presynaptic receptors that alter release. Paired-pulse ratio (PPR) was taken as the amplitude of axEPSP2/axEPSP1. For the shortest inter-stimulus interval of 10 ms, the PPR was calculated through subtraction (see methods). PPR was depressing for all interstimulus interval (ISI) shorter than 10 seconds (**Figure 5A,B**; PPR 0.11 ± 0.03 for 10 ms, n = 10 pairs; 0.27 ± 0.08 for 100 ms, n = 10; 0.54 ± 0.07 for 500 ms, n = 8; 0.78 ± 0.04 for 1000 ms, n = 10). In separate experiments, we tested PPR following optical stimulation of CINs. We observed no difference between the PPR curves for optically-evoked or electrically-stimulated axEPSPs suggesting that these results represent short-term plasticity of cholinergic transmission alone (**Figure 5A,B**; F(2, 76) = 1.28, p = 0.28; Tukey post-hoc test q(76) = 0.91, p = 0.79).

**Figure 5.**
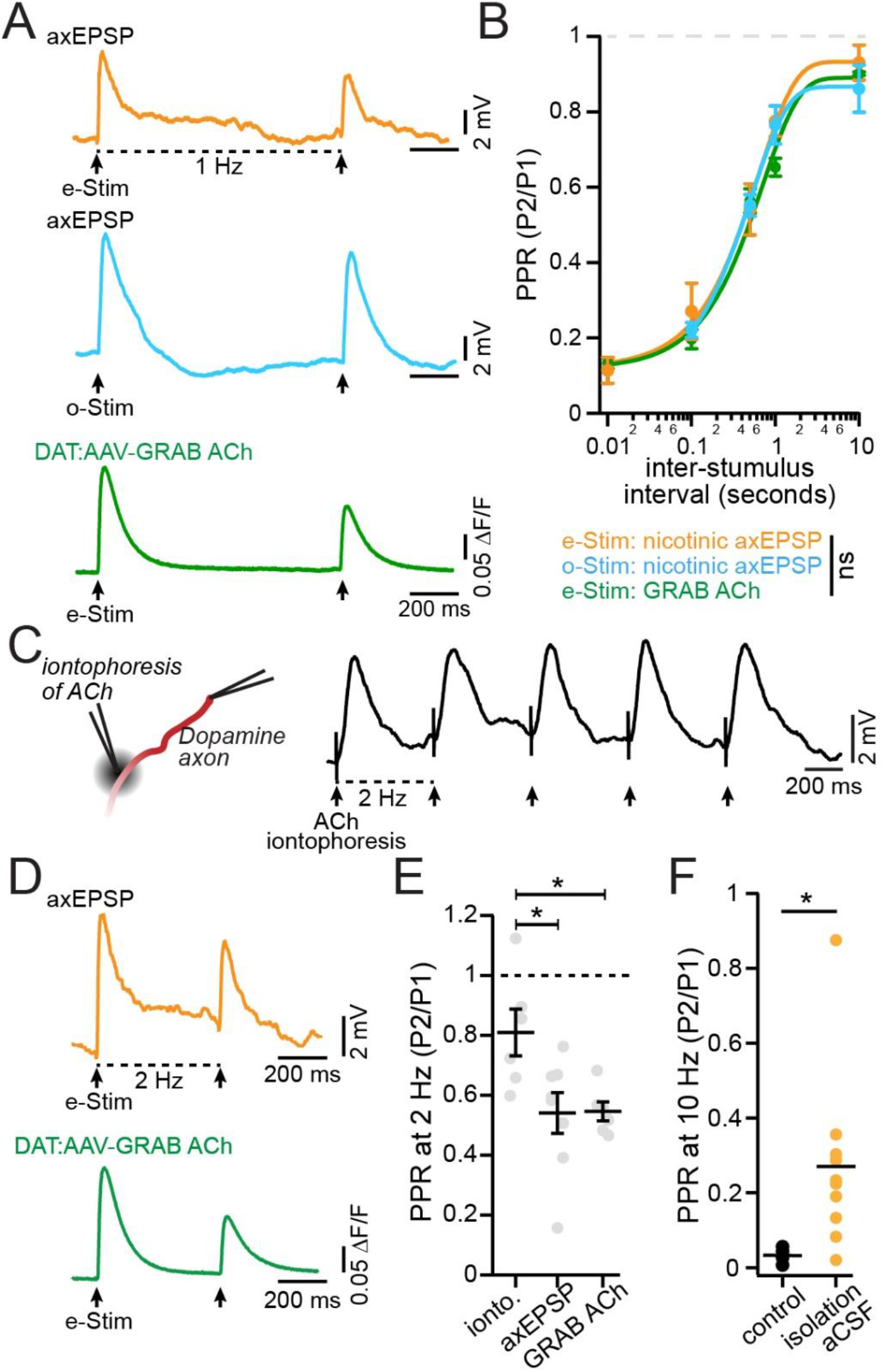
Desensitization of nAChRs contributes little to short-term plasticity. **A.** Example traces for 1 Hz paired-pulse stimulation of: electrically-evoked axEPSP in isolation aCSF (top row), optical activation of CINs to evoke an axEPSP (middle row), and electrical stimulation in isolation aCSF and recorded with the GRAB_ACh_ fluorescent ACh sensor (GACh). **B.** Combined data for the full paired-pulse ratio response curves for all three conditions in (**A**). **C.** Experimental setup for iontophoresis of ACh onto DA axons during an axonal recording (left), and an example iontophoresis experiment showing 2 Hz application of ACh (right). **D.** Example traces for 2 Hz paired-pulse stimulation of electrically stimulated axEPSP in isolation aCSF (orange), and electrical stimulation in isolation aCSF and recorded with GACh (green). **E.** Comparison of the paired-pulse ratio at 2 Hz between iontophoresis (abbreviated “ionto.”, n = 6 axons), stimulated axEPSP (n = 8 axons), and stimulated GRAB_ACh_ (n = 9 slices). **F.** Comparison of the paired-pulse ratio at 10 Hz between stimulated axEPSPs recorded in control (n = 4 axons) and isolation (n = 10 axons) aCSF. *p<0.05

One possibility is that desensitization of nAChRs resulting from repeated stimulation may have contributed to the PPR depression of axEPSPs. To assess the possible contribution of receptor desensitization to the short-term depression described above, we measured cholinergic transmission using the fluorescent ACh sensor GRAB_ACh_ (Jing et al., 2020) which we selectively expressed in dopamine neurons. We reasoned that if receptor desensitization contributes to the depression of nicotinic axEPSPs, then GRAB_ACh_ signals should show significantly less frequency-dependent depression. In contrast to this idea, however, we found no difference in PPR between nicotinic axEPSPs and electrically-evoked GRAB_ACh_ signals (F(2, 76) = 1.28, p = 0.28; Tukey post-hoc test q(76) = 2.26, p = 0.25; **Figure 5A,B**).

In complementary experiments, we tested directly whether repeated activation of nicotinic receptors produces desensitization. To test this, we recorded the axonal membrane potential in perforated-patch while exogenously applying ACh to dopamine axons using iontophoresis (**Figure 5C**). We found that iontophoresis pulses applied at 500 ms intervals resulted in significantly less depression relative to electrically-evoked nicotinic axEPSPs and stimulated GRAB_ACh_ signals at the same intervals (iontophoresis-PPR = 0.81 ± 0.08 at 500 ms; e-stim PPR = 0.54 ± 0.07 at 500 ms; GRAB_ACh_ PPR = 0.56 ± 0.02; ionto vs e-stim: Z(8) = 2.34, p = 0.038; ionto. vs GRAB_ACh_ Z(9) = 2.75, p = 0.012; **Figure 5D,E**). Finally, we observed a significantly larger depression of the 10 Hz stimulated axEPSP PPR when recordings were made in control relative to isolation aCSF (**Figure 5F**; control, PPR 0.03 ± 0.01 at 100 ms, n = 4 pairs; isolation aCSF, 0.27 ± 0.07 at 100 ms, n = 10 pairs; Mann-Whitney U = 3, p = 0.014), confirming a role for likely both presynaptic muscarinic and dopaminergic D2 receptors in inhibition of cholinergic transmission onto DA axons. Together, these data reveal short-term depression of CIN-mediated ACh release onto DA axons. Importantly, we demonstrate that this depression emerges from the properties of vesicular ACh release and not from nicotinic receptor desensitization, consistent with a previous study implicating vesicle depletion in mediating frequency-dependent depression (Wang et al., 2014). This finding is also consistent with past work finding less nicotinic receptor desensitization in dopamine neurons that express α6-containing nAChRs (Liu et al., 2012).

### Cholinesterase inhibition slows axEPSP decay kinetics but the amplitude is unaffected

We next tested the extent to which axo-axonal transmission between CIN and DA axons is regulated by AChE. The striatum has among the highest expression of AChE, an enzyme responsible for the hydrolysis of ACh, of anywhere in the brain (Hoover et al., 1978; Zhou *et al*., 2001). In other peripheral and central regions, AChE is localized to synapses where it is thought to limit free diffusion of ACh outside of the synaptic cleft following release events. To probe for possible receptors outside a synaptic contact, we used ambenonium to inhibit AChE, reasoning that this would have a predominant effect on receptors distal to release sites which require a longer diffusion distance (Bennett *et al*., 2012).

Inhibition of AChE using saturating concentration of ambenonium (300 nM or 1 µM) resulted in a brief increase in the frequency of spontaneous axEPSPs (159 ± 16.4% of control) followed by a long-lasting decrease (87.6 ± 12.8% of control, **Figure 6A-B**). Inhibition of AChE also resulted in shorter amplitude spontaneous axEPSPs, but only in the late phase of ambenonium treatment (**Figure 6C-D**; control: 1.00 ± 0.01 mV; amb-early: 1.02 ± 0.02 mV; amb-late: 0.87 ± 0.01 mV; ctrl-early p = 0.228; ctrl-late p = 0.005; n = 10). In addition, the kinetics of spontaneous axEPSPs were altered following inhibition of AChE, with a widening of the median half-width (**Figure 6E**; control, 54.0 ± 0.6 ms; amb-early 57.5 ± 0.82 ms; amb-late: 55.2 ± 1.26 ms; ctrl-early p = 0.0009; ctrl-late p < 0.0001) and a slowing of the median rise times (**Figure 6F**; control: 20.8 ± 0.23 ms; amb-early: 24.1 ± 0.28 ms; amb-late: 26.5 ± 0.33 ms; ctrl-early p = 0.003; ctrl-late p < 0.0001). Electrically-evoked axEPSPs were similarly affected by inhibition of AChE, as shown in the example traces in **Figure 6G**. The decay of the axEPSP was slowed substantially in ambenonium as reflected in a larger average area under the curve (AUC) across axons (**Figure 6I**; norm AUC in ambenonium: 1.98 ± 0.32, W = 28, p = 0.016, n = 7 axons). The axEPSPs also had longer rise times (**Figure6J;** control: 14.3 ± 2.48 ms; ambenonium: 26.5 ± 5.72 ms; Wilcoxon W=36, p = 0.008, n = 8 axons). These results are consistent with the idea that AChE inhibition produces longer lasting ACh signaling and broader ACh diffusion beyond CIN terminal release sites.

**Figure 6.**
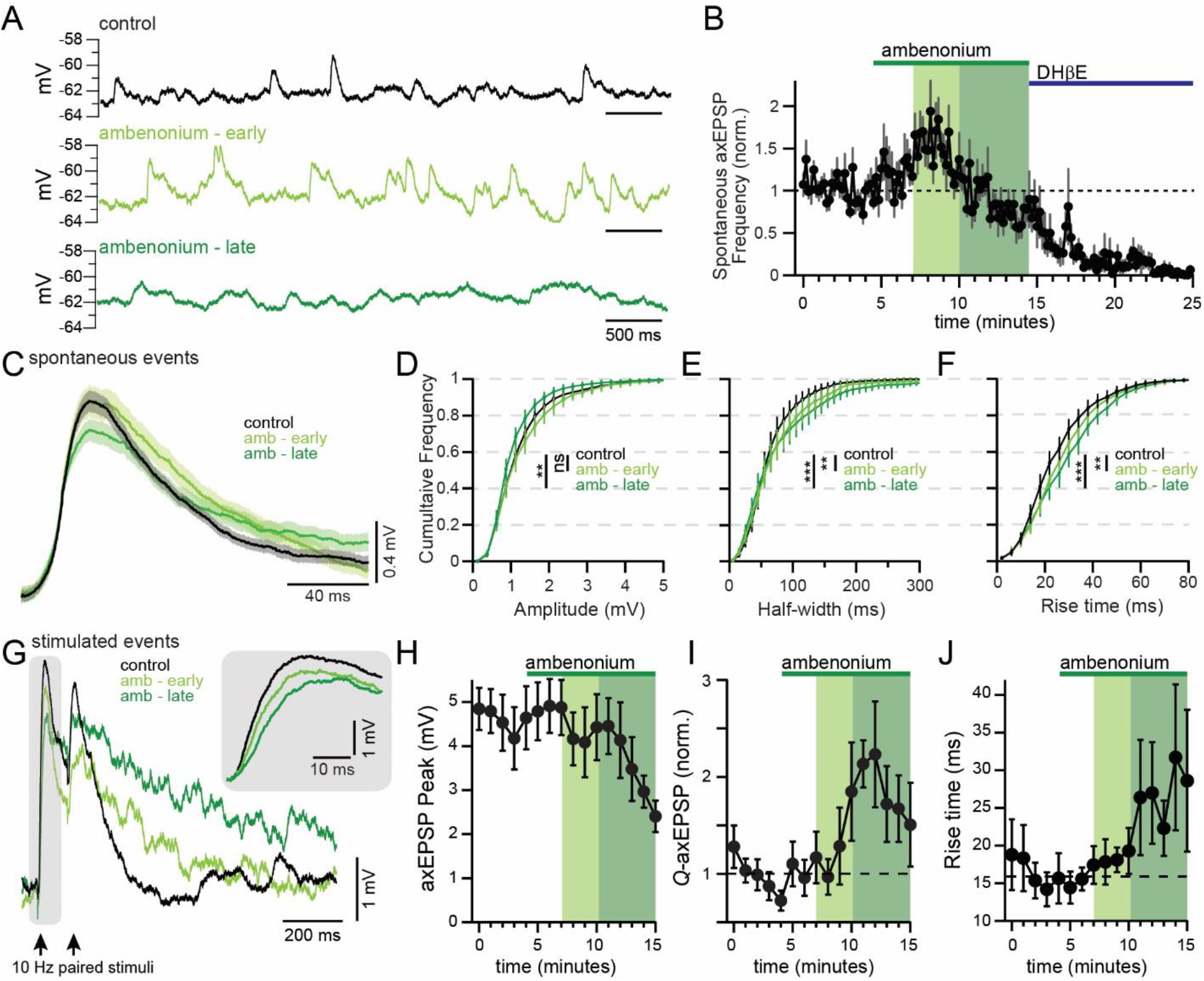
Kinetics of axEPSPs are shaped by AChE. **A.** Example traces of spontaneous axEPSPs in a dopaminergic axon in control (top), in the early phase of ambenonium wash-in (middle), and in the late phase of ambenonium (bottom). **B.** Time course showing the frequency of spontaneous axEPSPs (normalized) in response to the application of a saturating concentration of ambenonium (300 nM – 1 µM) followed by DHβE (1 µM). **C.** Example average (± s.e.m in shaded region) axEPSP from one axon in control (black) and in the early (light green) and late (dark green) phases of ambenonium. Cumulative histograms of axEPSP amplitude (**D**) half-width (**E**) and rise time (**F**) from all axons in control, early, and late phases of ambenonium. **G.** Example stimulated axEPSPs in control and in ambenonium. Inset shows an enhanced view of the rise time. Stimulated axEPSPs were analyzed for the effect of ambenonium on peak amplitude (**H**), area under the curve (**I**) and rise time (**J**), n = 8 axons. ***p < 0.001

Similar to analysis of volume transmission in other systems (Bennett *et al*., 2012; Coddington et al., 2013; Isaacson, 1999; Szapiro and Barbour, 2007), we reasoned that if diffusion of ACh across long distances is required to activate the nAChRs on DA axons then inhibition of AChE should result in an amplification of the axEPSP amplitude. Following inhibition of AChE, however, we found that ambenonium did not potentiate the peak amplitude of axEPSPs (**Figure 6D, G-H**). Rather, there was a small but significant reduction in peak amplitude from 4.8 ± 0.7 mV to 3.0 ± 0.3 mV in ambenonium (p = 0.031, n = 6 axons). Because CINs express presynaptic muscarinic autoreceptors that inhibit release, one possibility is that the reduction in peak axEPSP may result from autoreceptor inhibition caused by the increase in ambient ACh in ambenonium. However, recordings made in isolation aACSF that includes atropine to inhibit muscarinic receptors produced similar results (**Supplemental Figure 2**). This demonstrates that inhibition from muscarinic autoreceptors does not account for the lack of peak potentiation of axEPSPs in ambenonium. Together, these data show that AChE regulates the spatial and temporal release of ACh onto DA axons in the striatum. Unexpectedly, however, we found that inhibition of AChE does not result in potentiation of axEPSPs, a finding that is inconsistent with classical notions of volume transmission. These results therefore suggest an arrangement whereby a fraction of CIN terminals exist in close proximity to nAChRs on DA axons.

#### Nicotinic axEPSPs occur in DA axons of primates

To test the generality of our observations across species, we asked whether cholinergic transmission onto DA axons projecting to the putamen nucleus of non-human primates was also synaptic-like. We performed perforated-patch clamp recordings in DA axons of macaques (**Figure 7A-B**). We visualized dopaminergic fibers by incubating tissue slices in a fluorescently labeled cocaine analogue, MFZ9-18 (Eriksen et al., 2009), see methods. We observed spontaneous depolarizations as well as evoked axEPSPs, both of which were inhibited by DHβE (**Figure 7C-E**). We found the stimulated axEPSPs were similar in amplitude, kinetics, and short-term plasticity characteristics to stimulated axEPSPs recorded in mice (**Figure 7F-J**).

**Figure 7.**
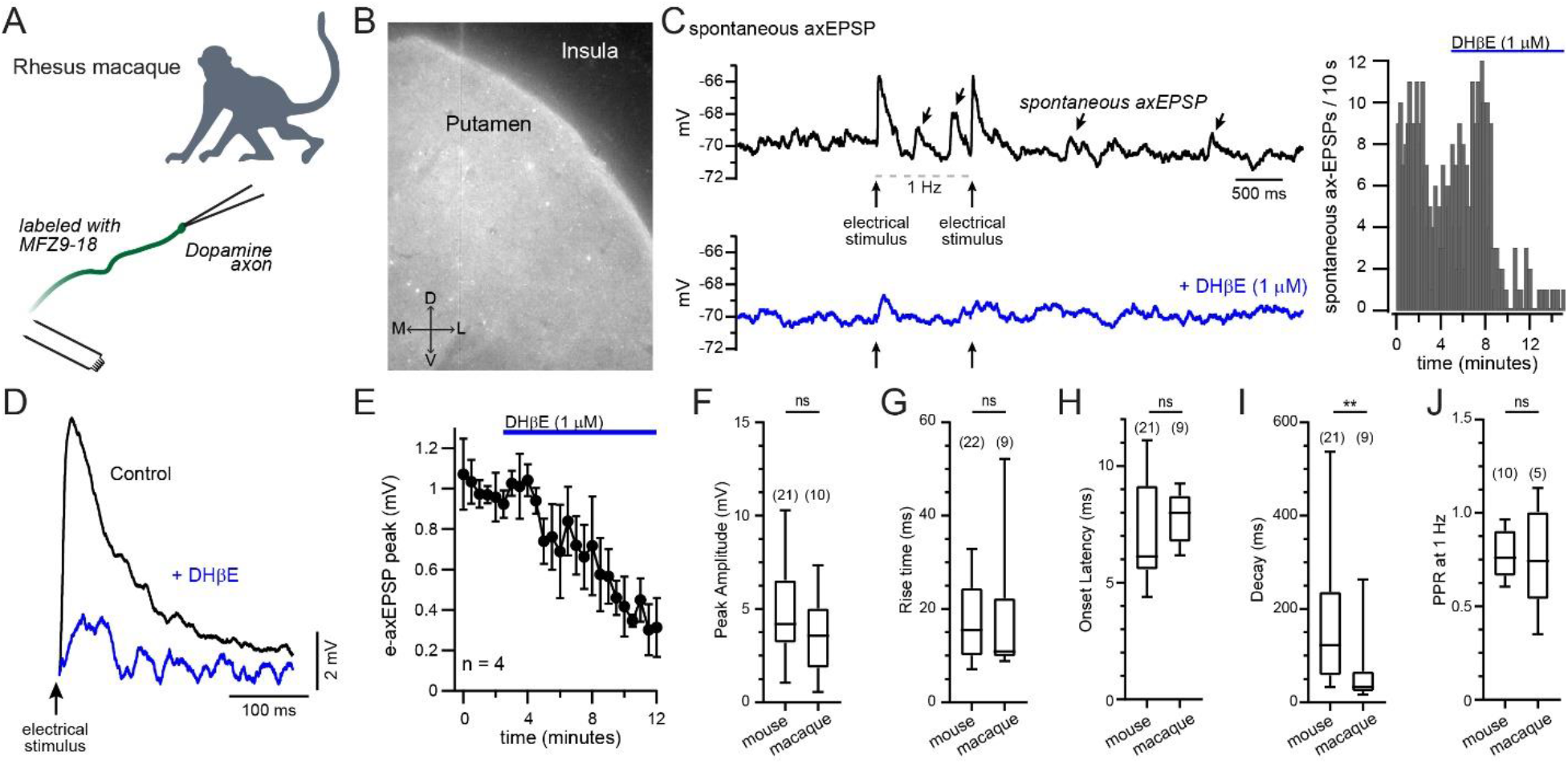
Spontaneous and evoked nicotinic axEPSPs recorded in DA axons of rhesus macaques. **A.** Experimental setup for experiments in rhesus macaque. DA axons were identified using MFZ9-18. A bipolar electrode was used for electrical stimulation. **B.** Picture of a macaque brain slice treated with MFZ9-18 to label DA axons. **C.** Example recording from a macaque dopaminergic axon showing spontaneous and stimulated axEPSPs in control (top, black) and after DHβE (bottom, blue). Right panel shows a histogram of the number of spontaneous axEPSPs for the axon in (**C**). **D.** Example traces of electrically stimulated axEPSPs in isolation aCSF (black) and following DHβE (blue) in macaque striatal brain slices. **E.** Time course showing the inhibition of the electrically stimulated axEPSP in macaque DA axons by DHβE (1 µM, n = 4 axons). The properties of the stimulated axEPSPs recorded in macaque were compared to those from the same conditions recorded in mice to analyze peak amplitude (**F**), rise time (**G**), onset latency (**H**), decay (**I,** **p<0.01), and paired-pulse ratio at 1 Hz (**J**). ns signifies p > 0.05.

Interestingly, the stimulated axEPSP decay was significantly faster in macaques (33.5 ± 26.1 ms, n = 9) than in mice (123 ± 33.3 ms, n = 21; U = 32, p = 0.004), indicating a potential difference in mechanisms regulating axEPSP decay such as AChE activity or channel properties. We also found that stimulated axEPSPs in macaque DA axons had a similar peak amplitude CV (0.23 ± 0.5) to those recorded in mice. Altogether, these results provide evidence that the synaptic-like depolarizations we record in mice translate to the DA axons of primates.

### AxEPSPs contribute to the generation of spontaneous APs in DA axon terminals

Studies from a variety of central neurons have proposed that the primary function of presynaptic nAChRs is to modulate neurotransmitter release, (Dani and Bertrand, 2007; Wonnacott, 1997) which can occur through several mechanisms. In the present study, we find that nAChR activity in DA axons by CINs likely modulates dopamine release by shaping the subthreshold voltage of DA axons. Yet, the presence of axEPSPs in DA axons raises the question of whether summation of axEPSPs may directly trigger APs in striatal DA axon terminals, independently of conventional somatic input and integration.

While most recorded axons lacked spiking activity as would be expected in recordings from isolated terminal arbors, we were surprised to find that a subset of axonal recordings (19/118 axons) exhibited nicotinic axEPSPs that were accompanied by spontaneous axonal APs (**Figure 8A, 8B**). Perforated-patch recordings with glass pipettes results in filtering of axonal APs (Olah et al., 2021; Ritzau-Jost et al., 2021). However, spontaneous APs could be identified as a characteristically abrupt increase in the first derivative of the membrane potential (d*V*/d*t*), which was also seen in all-or-none APs evoked from the same axons (**Figure 8C-E**). Furthermore, we compared spontaneous axonal APs recorded from the main trunk of the dopamine axon in perforated-patch recordings and constructed a weighted z-score to identify spontaneous APs in striatal perforated-patch recordings (**Supplemental Figure 3**). Using this method, we identified 63 spontaneous axonal APs recorded from 19 axonal recordings. We found that in 45 APs, the occurrence of the axonal spike overlapped with the depolarization from an axEPSP with the majority of spikes occurring during the rising phase (**Figure 8F and G**). The remaining 18 APs occurred in the absence of any resolvable axEPSP, suggesting active propagation of the AP past the point where the subthreshold axEPSP had passively decayed. These data strongly suggest that nicotinic release events that produce axEPSPs are capable of triggering spontaneous all-or- none APs in the terminal region of a DA axons.

**Figure 8.**
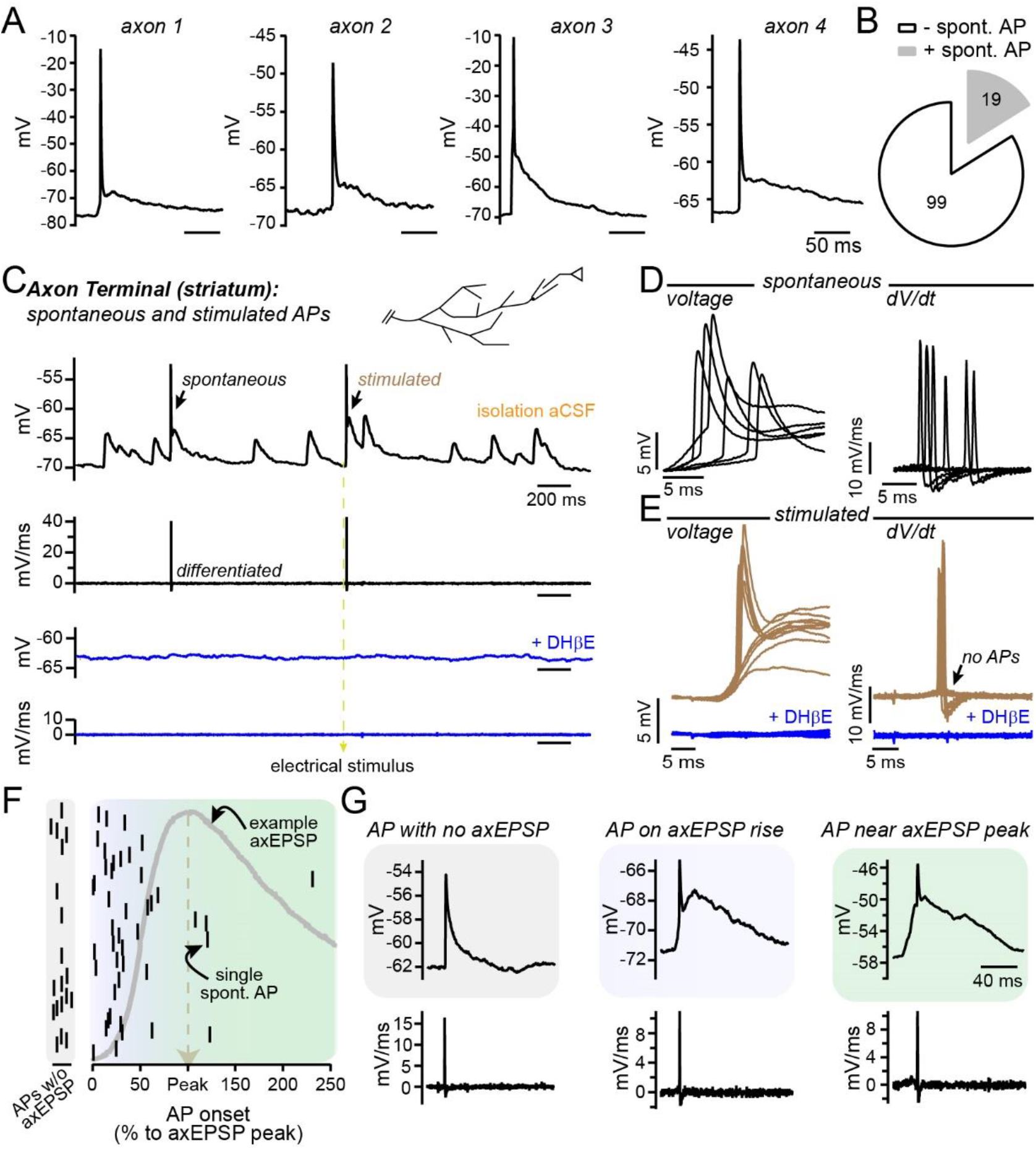
Spontaneous APs in DA axons triggered by nicotinic axEPSPs. **A.** Four example spontaneous APs recorded in four separate DA axons. **B.** Pie chart of the number of axonal recordings that did (grey) or did not (white) contain at least one positively identified spontaneous AP. **C.** Example recoding from a dopamine axon in isolation solution. Left arrow points to a spontaneous AP, right arrow points to an AP following an electrically stimulated axEPSP. Bottom black trace is the differentiation. Blue traces below are from the same time window as the control traces after DHβE application, showing voltage (top, blue) and differentiated trace (bottom, blue). **D.** All spontaneous APs from the same recording shown in (**C**), aligned to the onset of the axEPSP (voltage on the left, differentiation on right). **E.** All stimulated axEPSPs from the same recording shown in (**C**), aligned to 4 ms before stimulation (voltage on the left, differentiation on right), blue shows the same time window after DHβE. **F.** Plot of AP timing relative to its underlying axEPSP (APs with no axEPSP are separated to the left). The y-axis is a linear count. **G.** Three examples of spontaneous APs in three different scenarios: with no underlying axEPSP (left panel), occurring on the rising phase of the axEPSP (middle panel), or occurring near the axEPSP peak (right panel).

## DISCUSSION

In this study, we examine the mechanisms of cholinergic transmission onto DA axons and their control of axonal excitability. Our results demonstrate a hybrid model of cholinergic transmission onto DA axons that has non-synaptic properties but, surprisingly, also exhibits several functional hallmarks of signaling observed at standard point-to point synapses. We find that axonal EPSPs generated during endogenous activity of CINs are capable of triggering APs spontaneously in DA axon terminals. We establish that α6-containing nAChRs contribute significantly to axEPSP in DA axons. Our experiments also demonstrate that short-term depression of nAChR activation is not limited by nAChR desensitization but results from presynaptic release mechanisms of cholinergic interneurons. Finally, our recordings from DA axons in the macaque putamen show that this unique form of axo-axonal synaptic transmission is not limited to rodents but exists also in primates.

### Synaptic and non-synaptic functional consequences of ACh release on DA axons

Previous conceptualizations of cholinergic transmission onto DA axons were based solely on structural studies which predicted that presynaptic nAChRs are likely activated by diffuse volume transmission (Descarries et al., 1997). EM studies reported a low occurrence of synaptic specializations opposite striatal cholinergic boutons (Aznavour *et al*., 2003; Contant *et al*., 1996; Izzo and Bolam, 1988) and that β2-positive nAChRs on DA axons are only rarely apposed to presynaptic structures (Jones *et al*., 2001). One electron microscopy (EM) structural study in striatum found no identifiable synaptic connections between cholinergic terminals and DA axons but reported that tight appositions between tyrosine hydroxylase positive axons and ChAT positive axons were numerous, suggestive of potential interactions (Chang, 1988). Despite evidence that DA axons may be located near cholinergic terminals, however, the lack of synaptic specializations suggests that volume transmission may be the main mode of activation for nAChRs on DA axons.

In contrast to the slow signaling associated with volume transmission, our functional recordings suggest that cholinergic transmission onto DA axons occurs over a range of signaling speeds that includes rapid signaling through axonal nAChRs. Quantification of spontaneous and evoked axEPSPs revealed a distribution of fast rise times with the median value being at approximately 20 ms and a relatively wide standard deviation to include fast (<10 ms) and slow (>30 ms) events. These values are roughly comparable to those reported for fast cholinergic transmission onto somatodendritic postsynaptic sites in cells from retina, cortex, and hippocampus (Bennett *et al*., 2012; Obermayer *et al*., 2019; Sethuramanujam et al., 2021; Sun *et al*., 2013) and differ significantly from values of slow nicotinic transmission that can have rise times of up to 100 ms. It is important to consider that the kinetic values reported here are likely influenced by our experimental recording conditions. For example, the perforated-patch configuration can significantly filter rapid membrane depolarizations. In addition, the rise times of the EPSPs will be slowed due to temporal filtering from cable properties as they travel from release site through the thin DA axons to our recording electrodes. Therefore, it is likely that the axEPSP rise times reported in our study are an underestimate of cholinergic signaling speed at the axonal site of nAChR activation.

Our results show axEPSPs exhibit functional characteristics that are typically observed in conventional synaptic transmission. We observed spontaneous axEPSPs following block of AP-dependent transmission (i.e. miniature EPSPs), which indicates that a population of nAChRs on DA axons are activated following release from a single vesicle of ACh. Furthermore, we found that the peak amplitude of electrically-evoked axEPSPs was insensitive to AChE inhibition by ambenonium. This demonstrates that transmission onto nAChRs activated during the earliest axEPSP phase is not limited by AChE, suggesting that these receptors may be closely localized to release sites. Interestingly, the decay of axEPSPs was slowed substantially in response to AChE inhibition suggesting that the later phase of stimulated axEPSPs involves diffusion to distant nAChRs (Bennett *et al*., 2012). Consistent with this observation, we found that inhibition of AChE led to a transient increase in the frequency of detectable spontaneous axEPSPs likely resulting from activation of nAChRs at newly accessible sites at longer diffusion distances. Analyzing the peak amplitude of stimulated axEPSP, we see a relatively small coefficient of variation of our stimulated axEPSP of ∼0.16, a value which is comparable to non-synaptic glutamate transmission in the cerebellum and cortex (Bennett *et al*., 2012; Szapiro and Barbour, 2007). Finally, the latency to 5% of the peak axEPSP (onset) is ∼6 ms, which aligns closely with an estimate for cholinergic transmission onto dopamine axons extrapolated from carbon fiber recordings of dopamine release (Wang *et al*., 2014). This latency to onset is relatively slow compared to conventional synaptic transmission for glutamate (Geiger et al., 1997; Szapiro and Barbour, 2007) and ACh (Hefft et al., 1999). These data create a picture of cholinergic transmission onto DA axons as having functional properties consistent with synaptic transmission, but also likely includes signaling at non-synaptic sites. In light of past ultrastructural studies and our functional data, therefore, we propose a hybrid model of transmission that allows for volume-mediated release of ACh to produce subthreshold depolarizations of the axonal membrane potential that are fast and have both synaptic and non-synaptic characteristics. Such a model has recently been proposed to underlie cholinergic signaling in the brain, rejecting dichotomous conceptualization of neurotransmission as either synaptic or non-synaptic in favor of an appreciation for a nuanced continuum of transmission modalities (Disney and Higley, 2020).

### Towards understanding the biomechanics of axo-axonal CIN to DA transmission

How can lines of evidence that suggest both synaptic and non-synaptic signaling be incorporated into a unified mechanistic framework? One possibility is that the synaptic-like behavior results from a small number of close apposition sites. This could arise from directed formation of synaptic contacts, but the ultrastructural evidence does not support that hypothesis. As a non-exclusive alternative, there could be a random distribution of ACh release sites and DA axon nicotinic receptors that are in high enough density to allow for occasional synaptic-like contacts. In addition, the unique physiology of DA axons could make a traditional synaptic contact unnecessary for a functionally synaptic response. DA axons have a high input resistance of ∼1.8 GΩ (Kramer *et al*., 2020) and express nAChRs with a high affinity for ACh, particularly the α6-containing receptors which have an EC_50_ for ACh of approximately 100 nM and are the least well understood physiologically (Chen et al., 2018; Drenan et al., 2008; Grady et al., 2010; Gu et al., 2019; Salminen et al., 2007). These characteristics of the DA axon would allow for large nAChR signals from only a few receptors. Similar results have been observed in the retina where starburst amacrine cell-mediated release of ACh onto direction-sensitive ganglion cells is thought to occur at multiple post-synaptic sites. Here it was reported that release of ACh spans an area of hundreds of nanometers to a couple of micrometers, and still the currents recorded in the ganglion cells were largely synaptic in their functional properties (Sethuramanujam *et al*., 2021). In order to fully understand how these synaptic-like nicotinic EPSPs arise in the dopamine axon, a better understanding of subcellular spatial dynamics of ACh release and diffusion in the extracellular space around DA axons is needed.

Here, we report that the α6 subunit of the nAChR accounts for a majority of the evoked and spontaneous axEPSPs recorded in DA axons. Interestingly, the slowing of axEPSP rise times following Ctx-P1a suggests further diffusion to receptors following vesicular release of ACh, consistent also with the smaller amplitude events we observe. To explain this, one possibility is that α6-containing nAChRs may be positioned closer to ACh release sites relative to receptors lacking this subunit. Alternatively, these results could be explained by inherit differences in α4 versus α6 kinetics, a question that has not been well explored due to difficulties in expressing α6-containing nicotinic receptors in heterologous systems (Gu *et al*., 2019). Lastly, the higher affinity of α6-containing nicotinic receptors may enable the activation of nAChRs distal to the release site with reasonable efficiency. To differentiate between these hypotheses, the mechanics and pharmacodynamics of the α6 nicotinic receptors will need to be more fully explored in tightly controlled expression systems.

### Nicotinic signaling on axons from macaques

The role for striatal CIN activity in rewarding behaviors and neurodegenerative diseases has often been demonstrated through *in vivo* recordings in non-human primates (Aosaki *et al*., 1994; Apicella et al., 1997; Kimura et al., 1984; Raz et al., 1996). Yet, mechanistic research examining cholinergic modulation of dopamine release has largely been performed in rodent animal models. We found spontaneous and stimulated activation of nicotinic receptors on DA axons of rhesus macaques that largely resembled the events we recorded in mice in terms of kinetics, amplitude, and latency to onset. One interesting difference was in the stimulated axEPSP decay kinetics, which were significantly faster in macaque. This difference may reflect a sharper integration time window in macaques for nAChR-evoked axonal APs. Therefore, the generation of axonal spikes may require strong synchrony in cholinergic interneuron firing in primates.

### Independent input-output signaling of dopaminergic axons

The dendrites of dopamine neurons perform roles traditionally associated with axons, reliably backpropagating action potentials (Hage and Khaliq, 2015; Hausser et al., 1995) and releasing dopamine (Beckstead et al., 2004; Gantz et al., 2013; Gantz et al., 2018; Hikima et al., 2021; Lebowitz et al., 2021). In this study we reveal properties of DA axons that are reminiscent of dendritic functions. We report the presence of spontaneously occurring EPSPs that have traits similar to events recorded in the somatodendritic region. We show that these EPSPs generate spontaneous APs originating in the dopaminergic axon, suggestive of integrative processing independently occurring in the axonal compartment. In this scenario, transmission at nicotinic receptors is the excitatory input necessary for integration. In dendritic integration there are often inhibitory inputs that would balance out the excitatory drive. In the axon this inhibition may come from the shunting inhibition provided by axonal GABA-A receptors (Kramer *et al*., 2020). Interestingly, axonal GABA-A signaling lacks the rapid characteristics of axonal cholinergic transmission, perhaps setting a GABA-dependent permissive or restrictive tone for the generation of APs in the axon by cholinergic input.

The extent to which the axons of DA neurons act independently from their associated somatic signal during DA-dependent behaviors has been a topic of significant recent interest (Hamid et al., 2016; Kim et al., 2020; Mohebi *et al*., 2019). With our observation of axonal cholinergic EPSPs, we reveal a mechanism for how DA axons function beyond simple spike transmission to potentially include a role in signal processing, a function that has not been classically associated with axons. In particular, the Law of Dynamic Polarization has guided the field of neuroscience stating that information arrives in the dendrites, is integrated in the somas, and then travels down the axon to the next synapse (Ramon y Cajal, 1899). In this view, axons function to transmit signals but contribute little to signal processing and vice versa for dendrites. Recently, there is an increasingly detailed understanding of analog signaling in the axon and its role in output modulation (Rama et al., 2018). Here, we have taken the first look at endogenous subthreshold transmission onto a central axon, showing an unexpected contribution of synaptic-like cholinergic transmission that suggests an autonomous role for axons in neuronal processing. Presynaptic nAChRs are expressed throughout the brain, presenting an intriguing question of what other neurotransmitter systems may exhibit the type of axonal neurotransmission reported here, and what this form of transmission means for a broader understanding of how neural circuitry is anatomically and functionally organized.

## ACKNOWLEDGEMENTS

We thank Drs Veronica Alvarez, Jeffery Diamond, Bruce Bean, as well as members of the Khaliq laboratory for their insightful discussions and comments on this manuscript. This work was carried out in collaboration with the Non-Human Primate Physiology Consortium (NPPC) at the NIH Intramural Program. A portion of this study was funded by a fellowship to P.F.K. from the Center for Compulsive Behaviors, NIH Intramural Program. This work was supported by NINDS Intramural Research Program Grant NS003135 to Z.M.K., NIMH Intramural Research Program Grant MH002928 to B.B.A., and NIDA Intramural Research Program Grant DA000610 to A.H.K.

## KEY RESOURCES TABLE

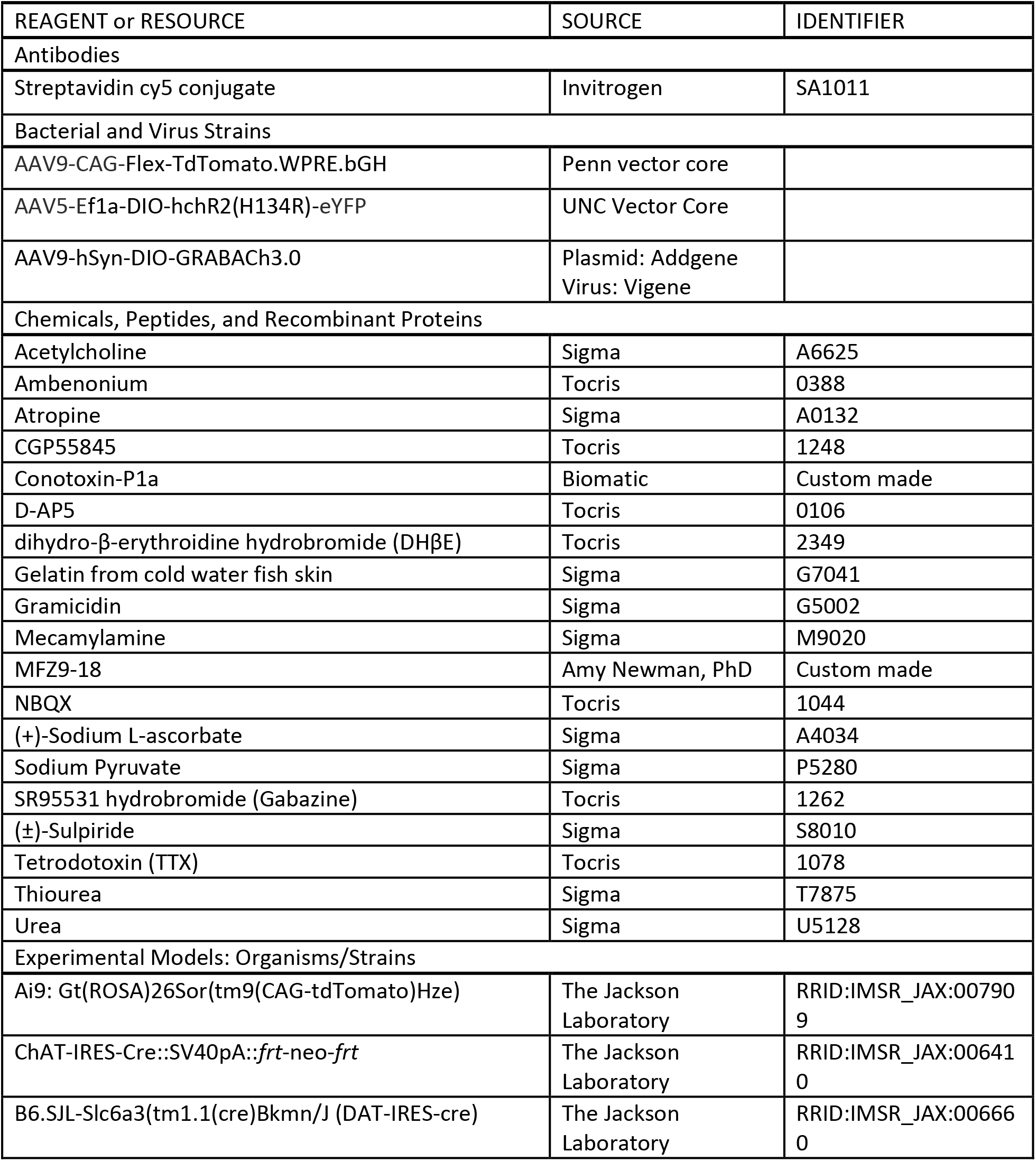

## STAR METHODS

### Resource Availability

#### Lead Contact and Materials Availability

Further information and requests for resources and reagents should be directed to and will be fulfilled by the Lead Contact Zayd Khaliq, Ph.D.

#### Data and Code Availability

- All data reported in this paper will be shared by the lead contact upon request.
- This paper does not report original code.
- Any additional information required to reanalyze the data reported in this paper is available from the lead author upon request.

### Experimental Model and Subject Details

#### Mice

All animal handling and procedures were approved by the animal care and use committee (ACUC) for the National Institute of Neurological Disorders and Stroke (NINDS) at the National Institutes of Health. Mice of both sexes were used throughout the study. Mice that underwent viral injections were injected at postnatal day 18 or older and were used for *ex vivo* electrophysiology and imaging 3-12 weeks after injection. The following strains were used: DAT-Cre (RRID:IMSR_JAX:006660); ChAT-cre (RRID:IMSR_JAX:006410); Ai9 (RRID:IMSR_JAX:007909).

#### Rhesus Macaques

All experimental procedures were performed in accordance with the ILAR Guide for the Care and Use of Laboratory Animals and were approved by the Animal Care and Use Committee of the National Institute of Mental Health. Procedures adhered to applicable United States federal and local laws, regulations, and standards, including the Animal Welfare Act and Regulations (PL89–544; 1985 https://www.nal.usda.gov/awic/animal-welfare-act) and Public Health Service (PHS) Policy (PHS2002). Two female rhesus macaques (*Macaca mulatta*, *P* - 12.1yo, PAG lesion and rear split; *MJ* - 9.8yo, bilateral amygdala excitotoxic lesions) were used for this study. For brain extraction, the animals were sedated with ketamine/midazolam (ketamine 5-15 mg/kg, midazolam 0.05-0.3 mg/kg) and maintained on isoflurane. A deep level of anesthesia was verified by an absence of response to toe-pinch and absence of response to corneal reflex. The animal was perfused with ice-cold artificial CSF solution containing in mM: 124 NaCl; 23 NaHCO_3_; 3 NaH_2_PO_4_; 5 KCl; 2 MgSO_4_; 10 d-glucose; 2 CaCl_2_.

### Method Details

#### Viral Injections

All stereotaxic injections were conducted on a Stoelting QSI (Cat#53311). Mice were maintained under anesthesia for the duration of the injection and allowed to recover from anesthesia on a warmed pad. Viruses used for this study were: AAV9-CAG-Flex-TdTomato.WPRE.bGH (Penn Vector Core; AV-9-ALL864); AAV9-hSyn-DIO-GRABACh3.0 titer: > 1 x 10^12^, plasmid: Addgene (#121923), virus packaged by Vigene; and AAV5-Ef1a-DIO- hchR2(H134R)-eYFP, titer: 4×10^12^, UNC vector core. Viral aliquots were injected (0.5-1 µl) bilaterally into either the medial dorsal striatum (X: ± 1.7 Y: +0.8 Z: -3.3) or the SNc (X: ± 1.9 Y: - 0.5 Z: -3.9) via a Hamilton syringe. At the end of the injection, the needle was raised at a rate of 0.1 to 0.2 mm per minute for 10 minutes before the needle was removed.

#### Slicing and electrophysiology

Brain slice experiments were performed on male and female adult mice of at least 6 weeks in age. Mice were anesthetized with isoflurane, decapitated, and brains rapidly extracted. Horizontal sections were cut at 330-400 µm thickness on a vibratome while immersed in warmed, modified, slicing ACSF containing (in mM) 198 glycerol, 2.5 KCl, 1.2 NaH_2_PO_4_, 20 HEPES, 25 NaHCO_3_,10 glucose, 10 MgCl_2_, 0.5 CaCl_2_. Cut sections were promptly removed from the slicing chamber and incubated for 30-60 minutes in a heated (34°C) chamber with holding solution containing (in mM) 92 NaCl, 30 NaHCO_3_, 1.2 NaH_2_PO_4_, 2.5 KCl, 35 glucose, 20 HEPES, 2 MgCl_2_, 2 CaCl_2_, 5 Na-ascorbate, 3 Na-pyruvate, and 2 thiourea. Slices were then stored at room temperature and used 30 min to 6 hours later. Following incubation, slices were moved to a heated (33–35°C) recording chamber that was continuously perfused with recording aCSF (in mM): 125 NaCl, 25 NaHCO_3_, 1.25 NaH_2_PO_4_, 3.5 KCl, 10 glucose, 1 MgCl_2_, 2 CaCl_2_.

To prepare macaque brain slices, tissue blocks were prepared containing the caudate and putamen and were then mounted on a vibratome as described for mice. Coronal sections were cut at 300 µm thickness in ice-cold sucrose cutting solution, filtered through a 0.22 µm filter (Nalgene), that contained (in mM): 90 sucrose, 80 NaCl, 3.5 KCl, 24 NaHCO_3_, 1.25 NaHPO_4_, 4.5 MgCl_2_, 0.5 MgCl_2_, 10 glucose. Cut section were promptly placed into a heated (34°C) chamber with the same sucrose solution and incubated in the warm bath for 30-60 minutes. Slices were then stored at room temperature and used 30 minutes to 3 days later. The incubation solution was exchanged for clean solution every 12 hours.

Perforated-patch recordings were made using borosilicate pipettes (5-10 MΩ) filled with internal solution containing (in mM) 135 KCl, 10 NaCl, 2 MgCl2, 10 HEPES, 0.5 EGTA, 0.1 CaCl2, adjusted to a pH value of 7.43 with KOH, 278 mOsm. Pipette tips were back filled with ∼1 µL of clean internal. Pipettes were then filled with internal containing between 80 and 100 µg/mL gramicidin. Patch integrity was monitored by the addition of either Alexa-488 (0.01 mM, axons filled with tdTomato) or Alexa-594 (0.01 mM, axons labeled with MFZ9-18) to the gramicidin-containing internal. Experiments where the perforation was lost were included up until the point where the recording substantially changed in quality (depolarized to above -45 mV, or changed input resistance by more than 15%). To enable post-hoc reconstruction, pipette solutions in some experiments included 0.1-0.3% w/v neurobiotin (Vector Labs). Current clamp recordings were manually bridge balanced. Conotoxin-P1a was synthesized based on the published structure by Biomatik (Ontario, CA).

Electrical stimulation was evoked with tungsten bipolar electrodes (150 µm tip separation, MicroProbes). For experiments where the site of electrical stimulation is distal to the site of imaging or recording, electrodes were placed at the caudal end of horizontal brain slices. Stimulations were evoked using an Isoflex (A.M.P.I.), amplitudes ranging from 0.8 to 20 V. ACh was rapidly applied near axons using iontophoresis with an MVCS-02C (NPI Electronic). ACh (1M) was loaded into a sharp micropipette (borosilicate glass, 40-60 MΩ), and the capacitance was corrected before each experiment. Iontophoretic pipettes were loaded with Alexa-488 (0.01 mM) to verify drug ejection during the experiment. During the experiment, square pulses (5-80 ms width) were applied to the pipette in a paired pulse (500 ms inter-stimulus interval) every 30 seconds. Atropine (100 nM) was included in the bath during iontophoresis experiments to prevent activation of mAChRs.

#### Fluorescent imaging

A white light LED (Thorlabs; SOLIS-3C) was used in combination with a tdTomato (Thorlabs; TLV-U-MF2-TOM) or EGFP (Chroma; 49002) filter set to visualize the DA axons (tdTomato) or for use with the ChR2 and GRAB_ACh_ (GFP). For visualizing the GRAB_ACh_ signals, a photodiode (New Focus) was mounted on the top port of the Olympus BX-51WI. For opsin activation, the LED was controlled with a TTL pulse (2-5 ms).

For visualization of DA axons in brain slices containing rhesus macaque tissue, a brain slice was moved to a separate incubation chamber with the regular sucrose solution, with the addition of 100 nM MFZ9-18 and saturating sulpiride (10 µM) for between 10 and 60 minutes. Slices were then moved to a recording chamber and axons visualized with blue light (EGFP filter set). MFZ9-18 was newly synthesized by Gisela Andrea Camacho-Hernandez, Ph.D. in the lab of Amy Newman, Ph.D. according to previously published protocols (Eriksen *et al*., 2009).

#### Immunohistochemistry, clearing, confocal imaging, and neural reconstructions

After electrophysiology or imaging, slices with neurobiotin filled axons were fixed overnight in 4% paraformaldehyde (PFA) in phosphate buffer (PB, 0.1M, pH 7.6). Slices were subsequently stored in PB until immunostaining and cleared using a modified CUBIC protocol, chosen because it does not quench endogenous fluorescence (Susaki et al., 2015). For the immunostaining and CUBIC clearing, all steps were performed at room temperature on a shaker plate. Slices were placed in CUBIC reagent 1 for 1-2 days, washed in PB 3 x 1 hour each, placed in blocking solution (0.5% fish gelatin (Sigma) in PB) for 3 hours. Slices were directly placed in streptavidin-Cy5 conjugate at a concentration of 1:1000 in PB for 2-3 days. Slices were washed 3 times for 2 hours each and were then placed in CUBIC reagent 2 overnight. Slices were mounted on slides in reagent 2 in frame-seal incubation chambers (Bio-Rad SLF0601) and coverslipped (#2 glass). Slices were imaged through 20×, 0.8 nA and 5×, 0.3 nA objectives on an LSM 800 confocal microscope (Zeiss) and taken as tiled z-stacks using Zen Blue software in the NINDS light imaging facility. Striatal axons were reconstructed using Neurolucida (MBF bioscience).

#### Quantification and Statistical Analysis

Fast voltages recorded from small compartments such as axons can appear distorted, especially when the recording is performed with perforated-patch which puts limitations on lowering series resistance (Olah *et al*., 2021). For this reason, putative spontaneous APs recorded for the analysis in **Figure 8** were compared to known APs stimulated in striatal perforated-patch recordings and to soma-generated spontaneous APs recorded in perforated-patch from the main trunk of the DA axon (Kramer *et al*., 2020). First, recordings were differentiated and signals above 10 standard deviations of the mean were selected as likely APs. A weighted z-score method was then used to establish which putative spontaneous APs to exclude from analysis that considered the absolute positive (*P)* and negative (*N*) peak d*V*/d*t* values, as well as the ratio 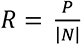 and the time *T = τ(N) – τ(P)*, between the two peak values. The time difference *T* was also used as a measurement of the AP half-width. These four values were established for the reference APs, and the putative spontaneous APs were compared against the averages using a z-score.

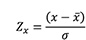

The z-scores were then combined using a weighted sum of squares to give weight to the relative values, which are more reliable to compare between experiments when doing axonal perforated-patch recordings (Olah *et al*., 2021).

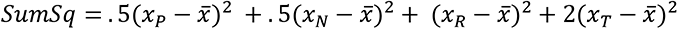

Putative APs with *SumSq* values larger than 21 (z-score of 3 for *P* and *N*, z-score of 2 for *R* and *T*) were excluded.

Spontaneous axEPSPs were detected using Neuromatic (Rothman and Silver, 2018), written for use with Igor Pro (Wavemetrics). We used event detection in Neuromatic that is based on a simple level detection algorithm (Kudoh and Taguchi, 2002). Levels were set at least 3 times the amplitude of baseline recording noise (between 0.3 and 1 mV), a sliding baseline average (5 ms) was used along with onset (1 SD, 30 ms) and peak detection (1 to 2 SD, 60 ms) parameters. After automated detection, traces were scanned for false positives, as defined by events without a clear decay following a rapid rise, and any such false positives were eliminated. Undetected events were only rarely manually added when automatic detection did not label a clear and obvious event, this was in order to not bias the data. See **Supplemental Figure 1** for three examples of automated peak detection output from three different axonal recordings. Spontaneous axEPSP kinetics were analyzed using built-in features in Neuromatic. Rise times were calculated as the time from 10 to 90% of the peak axEPSP amplitude. Decay was calculated as the time to 36.7% of the peak amplitude (1 time constant). Half-width was calculated as the time between the rise and decay points at half the peak axEPSP amplitude. Latency to onset was calculated as time between stimulus onset and 5% of peak axEPSP amplitude.

To calculate the paired-pulse ratio for the 10 ms inter-pulse intervals (100 Hz), the amplitude of the first pulse from the previous three traces was averaged and then subtracted from the amplitude of the 100 Hz train. The averaged amplitude was used as the pulse 1 value, and the subtracted amplitude was used as the pulse 2 values for P2/P1. A single exponential was used to fit the paired-pulse data.

Analysis was conducted in Igor Pro and Prism 8 (GraphPad). Data in text is reported in terms of means for parametric data and medians for non-parametric data. Error bars on graphs are indicated as ± SEM. Box plots show medians, 25 and 75% (boxes) percentiles, and maximum and minimum data points (whiskers). For parametric data, data in the text were expressed as means, t-tests were used for two-group comparison, and ANOVA tests were used for more than two group comparisons, followed by a Bonferonni or Tukey post-hoc test for analysis of multiple comparisons. For non-parametric data sets, data in the text was expressed as medians, Mann-Whitney U tests were used to compare two groups without repeated measures and a Wilcoxon test was used to compare two groups with repeated measures. A Kruskal-Wallis test was used to compare more than two groups followed by Dunn’s multiple comparison test for between-subject comparisons, while a Friedman test was used for comparisons within group (repeated measures) and a Dunn’s test used for multiple comparisons.

---

**Supplemental Figure 1.**
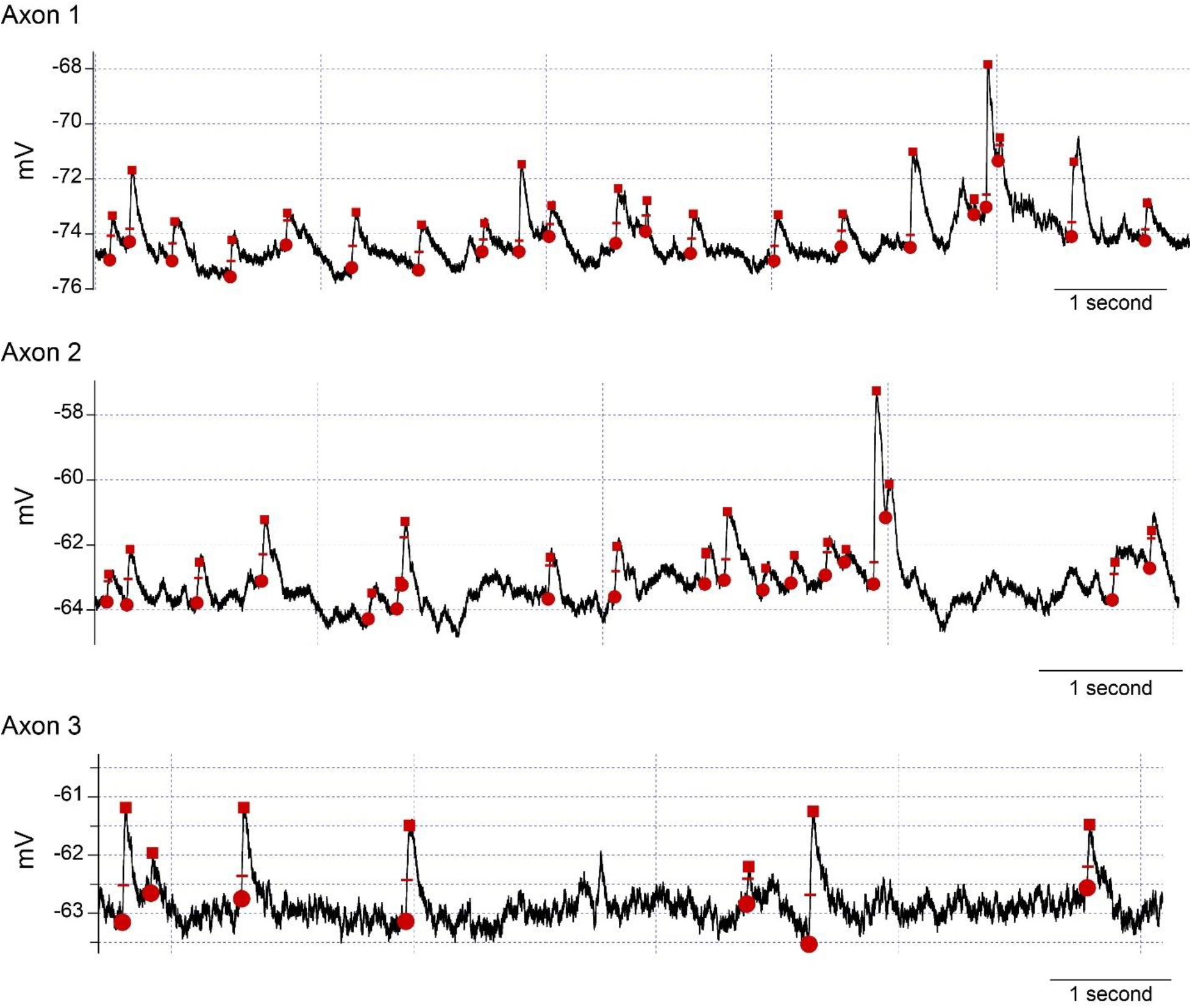
Example traces from three axonal recordings showing results from the automated detection algorithm used to detect spontaneous events. Axon 1 is shown in **Figure 1C**, axon 2 is shown in **Figure 6A**, and axon 3 is shown in **Figure 2A**.

**Supplemental Figure 2.**
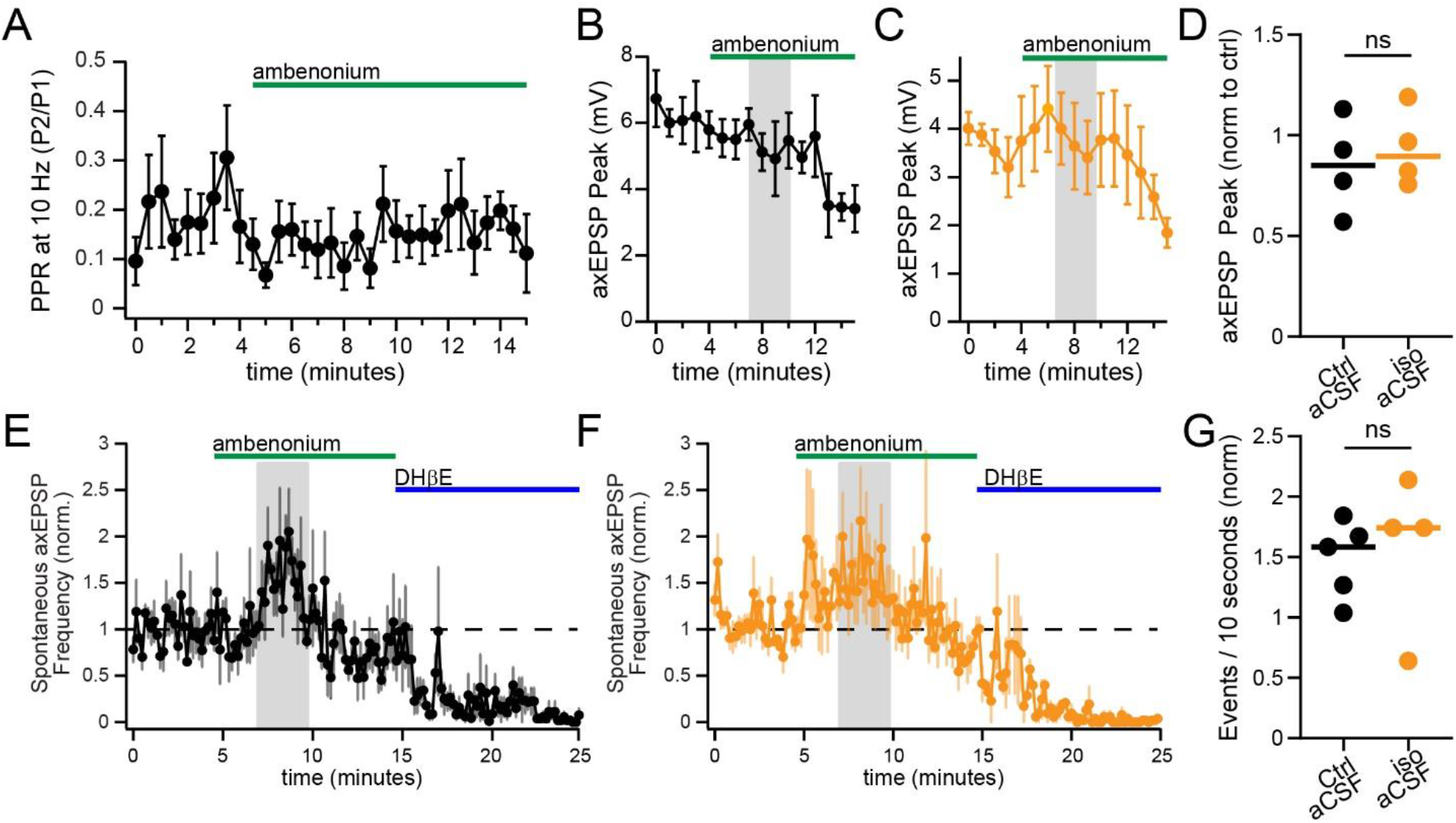
Expanded analysis of ambenonium. **A.** Lack of an effect of ambenonium on the 10 Hz axEPSP paired-pulse ratio (n = 6 axons). Comparison between control aCSF (**B**) and isolation aCSF (**C**) in the amplitude of the axEPSP peak during ambenonium wash-in. The early phase of ambenonium (2 to 5 minutes after wash-in begins) is averaged for each trace and plotted in **D** (control = 0.85 ± 0.12; iso = 0.93 ± 0.10; t(6) = 0.55, p = 0.60, n = 4 in each condition). Comparison between control aCSF (**E**) and isolation aCSF (**F**) in the frequency of spontaneous axEPSPs during ambenonium wash-in. The early phase of ambenonium (2 to 5 minutes after wash-in begins) is averaged for each trace and plotted in **G** (control = 1.48 ± 0.14, n = 5; iso = 1.57 ± 0.32, n = 4; t(7) = 0.26, p = 0.80). ns signifies p > 0.05.

**Supplemental Figure 3.**
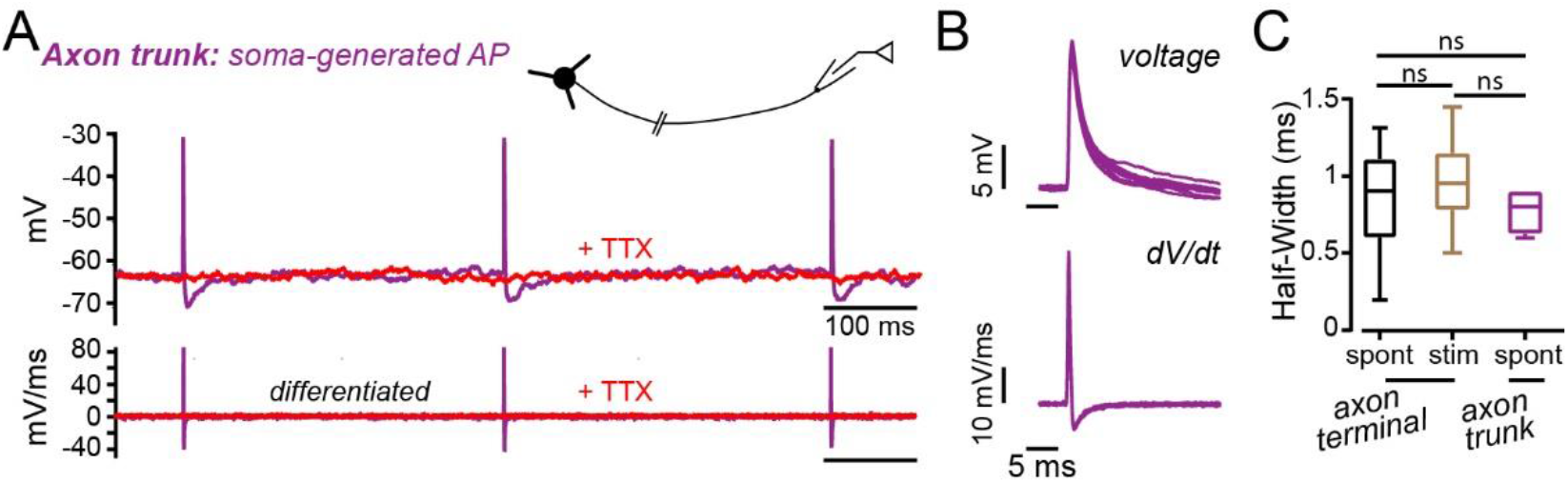
**A.** Example perforated-patch recording from the DA main axon trunk, showing spontaneously generated APs originating in the soma before (purple) and after (red) the application of TTX. Bottom shows the differentiated traces. **B.** Overlaid spontaneous APs from the recordings in (**A**), with voltage on the top and differentiated traces below. **C.** AP half-widths for spontaneous (spont) and stimulated (stim) APs recorded in striatal DA axons, as well as spontaneous APs recorded in the main axon trunk.

## References

1. Adrover, M.F., Shin, J.H., Quiroz, C., Ferre, S., Lemos, J.C., and Alvarez, V.A. (2020). Prefrontal Cortex-Driven Dopamine Signals in the Striatum Show Unique Spatial and Pharmacological Properties. J Neurosci 40, 7510–7522. 10.1523/JNEUROSCI.1327-20.2020.

2. Aosaki, T., Tsubokawa, H., Ishida, A., Watanabe, K., Graybiel, A.M., and Kimura, M. (1994). Responses of tonically active neurons in the primate’s striatum undergo systematic changes during behavioral sensorimotor conditioning. J Neurosci 14, 3969–3984.

3. Apicella, P., Legallet, E., and Trouche, E. (1997). Responses of tonically discharging neurons in the monkey striatum to primary rewards delivered during different behavioral states. Exp Brain Res 116, 456–466. 10.1007/pl00005773.

4. Aznavour, N., Mechawar, N., Watkins, K.C., and Descarries, L. (2003). Fine structural features of the acetylcholine innervation in the developing neostriatum of rat. J Comp Neurol 460, 280–291. 10.1002/cne.10660.

5. Beckstead, M.J., Grandy, D.K., Wickman, K., and Williams, J.T. (2004). Vesicular dopamine release elicits an inhibitory postsynaptic current in midbrain dopamine neurons. Neuron 42, 939–946. 10.1016/j.neuron.2004.05.019.

6. Bennett, C., Arroyo, S., Berns, D., and Hestrin, S. (2012). Mechanisms generating dual-component nicotinic EPSCs in cortical interneurons. J Neurosci 32, 17287–17296. 10.1523/JNEUROSCI.3565-12.2012.

7. Brimblecombe, K.R., Threlfell, S., Dautan, D., Kosillo, P., Mena-Segovia, J., and Cragg, S.J. (2018). Targeted Activation of Cholinergic Interneurons Accounts for the Modulation of Dopamine by Striatal Nicotinic Receptors. eNeuro 5. 10.1523/ENEURO.0397-17.2018.

8. Cachope, R., Mateo, Y., Mathur, B.N., Irving, J., Wang, H.L., Morales, M., Lovinger, D.M., and Cheer, J.F. (2012). Selective activation of cholinergic interneurons enhances accumbal phasic dopamine release: setting the tone for reward processing. Cell Rep 2, 33–41. 10.1016/j.celrep.2012.05.011.

9. Champtiaux, N., Gotti, C., Cordero-Erausquin, M., David, D.J., Przybylski, C., Lena, C., Clementi, F., Moretti, M., Rossi, F.M., Le Novere, N., et al. (2003). Subunit composition of functional nicotinic receptors in dopaminergic neurons investigated with knock-out mice. J Neurosci 23, 7820–7829.

10. Chang, H.T. (1988). Dopamine-acetylcholine interaction in the rat striatum: a dual-labeling immunocytochemical study. Brain Res Bull 21, 295–304. 10.1016/0361-9230(88)90244-4.

11. Chen, D.J., Gao, F.F., Ma, X.K., Shi, G.G., Huang, Y.B., Su, Q.X., Sudweeks, S., Gao, M., Dharshaun, T., Eaton, J.B., et al. (2018). Pharmacological and functional comparisons of alpha6/alpha3beta2beta3- nAChRs and alpha4beta2-nAChRs heterologously expressed in the human epithelial SH-EP1 cell line. Acta Pharmacol Sin 39, 1571–1581. 10.1038/aps.2017.209.

12. Coddington, L.T., Rudolph, S., Vande Lune, P., Overstreet-Wadiche, L., and Wadiche, J.I. (2013). Spillover-mediated feedforward inhibition functionally segregates interneuron activity. Neuron 78, 1050–1062. 10.1016/j.neuron.2013.04.019.

13. Collins, A.L., Aitken, T.J., Greenfield, V.Y., Ostlund, S.B., and Wassum, K.M. (2016). Nucleus Accumbens Acetylcholine Receptors Modulate Dopamine and Motivation. Neuropsychopharmacology 41, 2830–2838. 10.1038/npp.2016.81.

14. Contant, C., Umbriaco, D., Garcia, S., Watkins, K.C., and Descarries, L. (1996). Ultrastructural characterization of the acetylcholine innervation in adult rat neostriatum. Neuroscience 71, 937–947. 10.1016/0306-4522(95)00507-2.

15. Dani, J.A., and Bertrand, D. (2007). Nicotinic acetylcholine receptors and nicotinic cholinergic mechanisms of the central nervous system. Annu Rev Pharmacol Toxicol 47, 699–729. 10.1146/annurev.pharmtox.47.120505.105214.

16. Descarries, L., Gisiger, V., and Steriade, M. (1997). Diffuse transmission by acetylcholine in the CNS. Prog Neurobiol 53, 603–625. 10.1016/s0301-0082(97)00050-6.

17. Ding, J.B., Guzman, J.N., Peterson, J.D., Goldberg, J.A., and Surmeier, D.J. (2010). Thalamic gating of corticostriatal signaling by cholinergic interneurons. Neuron 67, 294–307. 10.1016/j.neuron.2010.06.017.

18. Disney, A.A., and Higley, M.J. (2020). Diverse Spatiotemporal Scales of Cholinergic Signaling in the Neocortex. J Neurosci 40, 720–725. 10.1523/JNEUROSCI.1306-19.2019.

19. Dowell, C., Olivera, B.M., Garrett, J.E., Staheli, S.T., Watkins, M., Kuryatov, A., Yoshikami, D., Lindstrom, J.M., and McIntosh, J.M. (2003). Alpha-conotoxin PIA is selective for alpha6 subunit-containing nicotinic acetylcholine receptors. J Neurosci 23, 8445–8452.

20. Drenan, R.M., Grady, S.R., Whiteaker, P., McClure-Begley, T., McKinney, S., Miwa, J.M., Bupp, S., Heintz, N., McIntosh, J.M., Bencherif, M., et al. (2008). In vivo activation of midbrain dopamine neurons via sensitized, high-affinity alpha 6 nicotinic acetylcholine receptors. Neuron 60, 123–136. 10.1016/j.neuron.2008.09.009.

21. Eriksen, J., Rasmussen, S.G., Rasmussen, T.N., Vaegter, C.B., Cha, J.H., Zou, M.F., Newman, A.H., and Gether, U. (2009). Visualization of dopamine transporter trafficking in live neurons by use of fluorescent cocaine analogs. J Neurosci 29, 6794–6808. 10.1523/JNEUROSCI.4177-08.2009.

22. Exley, R., Clements, M.A., Hartung, H., McIntosh, J.M., and Cragg, S.J. (2008). Alpha6-containing nicotinic acetylcholine receptors dominate the nicotine control of dopamine neurotransmission in nucleus accumbens. Neuropsychopharmacology 33, 2158–2166. 10.1038/sj.npp.1301617.

23. Gantz, S.C., Bunzow, J.R., and Williams, J.T. (2013). Spontaneous inhibitory synaptic currents mediated by a G protein-coupled receptor. Neuron 78, 807–812. 10.1016/j.neuron.2013.04.013.

24. Gantz, S.C., Ford, C.P., Morikawa, H., and Williams, J.T. (2018). The Evolving Understanding of Dopamine Neurons in the Substantia Nigra and Ventral Tegmental Area. Annu Rev Physiol 80, 219–241. 10.1146/annurev-physiol-021317-121615.

25. Geiger, J.R., Lubke, J., Roth, A., Frotscher, M., and Jonas, P. (1997). Submillisecond AMPA receptor- mediated signaling at a principal neuron-interneuron synapse. Neuron 18, 1009–1023. 10.1016/s0896-6273(00)80339-6.

26. Gopalakrishnan, M., Monteggia, L.M., Anderson, D.J., Molinari, E.J., Piattoni-Kaplan, M., Donnelly- Roberts, D., Arneric, S.P., and Sullivan, J.P. (1996). Stable expression, pharmacologic properties and regulation of the human neuronal nicotinic acetylcholine alpha 4 beta 2 receptor. J Pharmacol Exp Ther 276, 289–297.

27. Gotti, C., Guiducci, S., Tedesco, V., Corbioli, S., Zanetti, L., Moretti, M., Zanardi, A., Rimondini, R., Mugnaini, M., Clementi, F., et al. (2010). Nicotinic acetylcholine receptors in the mesolimbic pathway: primary role of ventral tegmental area alpha6beta2* receptors in mediating systemic nicotine effects on dopamine release, locomotion, and reinforcement. J Neurosci 30, 5311–5325. 10.1523/JNEUROSCI.5095-09.2010.

28. Grady, S.R., Drenan, R.M., Breining, S.R., Yohannes, D., Wageman, C.R., Fedorov, N.B., McKinney, S., Whiteaker, P., Bencherif, M., Lester, H.A., and Marks, M.J. (2010). Structural differences determine the relative selectivity of nicotinic compounds for native alpha 4 beta 2*-, alpha 6 beta 2*-, alpha 3 beta 4*- and alpha 7-nicotine acetylcholine receptors. Neuropharmacology 58, 1054–1066. 10.1016/j.neuropharm.2010.01.013.

29. Gu, S., Matta, J.A., Davini, W.B., Dawe, G.B., Lord, B., and Bredt, D.S. (2019). alpha6-Containing Nicotinic Acetylcholine Receptor Reconstitution Involves Mechanistically Distinct Accessory Components. Cell Rep 26, 866–874 e863. 10.1016/j.celrep.2018.12.103.

30. Hage, T.A., and Khaliq, Z.M. (2015). Tonic firing rate controls dendritic Ca2+ signaling and synaptic gain in substantia nigra dopamine neurons. J Neurosci 35, 5823–5836. 10.1523/JNEUROSCI.3904-14.2015.

31. Hamid, A.A., Pettibone, J.R., Mabrouk, O.S., Hetrick, V.L., Schmidt, R., Vander Weele, C.M., Kennedy, R.T., Aragona, B.J., and Berke, J.D. (2016). Mesolimbic dopamine signals the value of work. Nat Neurosci 19, 117–126. 10.1038/nn.4173.

32. Harvey, S.C., Maddox, F.N., and Luetje, C.W. (1996). Multiple determinants of dihydro-beta-erythroidine sensitivity on rat neuronal nicotinic receptor alpha subunits. J Neurochem 67, 1953–1959. 10.1046/j.1471-4159.1996.67051953.x.

33. Hausser, M., Stuart, G., Racca, C., and Sakmann, B. (1995). Axonal initiation and active dendritic propagation of action potentials in substantia nigra neurons. Neuron 15, 637–647. 10.1016/0896-6273(95)90152-3.

34. Hefft, S., Hulo, S., Bertrand, D., and Muller, D. (1999). Synaptic transmission at nicotinic acetylcholine receptors in rat hippocampal organotypic cultures and slices. J Physiol 515 (*Pt 3*), 769–776. 10.1111/j.1469-7793.1999.769ab.x.

35. Hikima, T., Lee, C.R., Witkovsky, P., Chesler, J., Ichtchenko, K., and Rice, M.E. (2021). Activity-dependent somatodendritic dopamine release in the substantia nigra autoinhibits the releasing neuron. Cell Rep 35, 108951. 10.1016/j.celrep.2021.108951.

36. Hoover, D.B., Muth, E.A., and Jacobowitz, D.M. (1978). A mapping of the distribution of acetycholine, choline acetyltransferase and acetylcholinesterase in discrete areas of rat brain. Brain Res 153, 295–306. 10.1016/0006-8993(78)90408-0.

37. Howe, M., Ridouh, I., Allegra Mascaro, A.L., Larios, A., Azcorra, M., and Dombeck, D.A. (2019). Coordination of rapid cholinergic and dopaminergic signaling in striatum during spontaneous movement. Elife 8. 10.7554/eLife.44903.

38. Isaacson, J.S. (1999). Glutamate spillover mediates excitatory transmission in the rat olfactory bulb. Neuron 23, 377–384. 10.1016/s0896-6273(00)80787-4.

39. Izzo, P.N., and Bolam, J.P. (1988). Cholinergic synaptic input to different parts of spiny striatonigral neurons in the rat. J Comp Neurol 269, 219–234. 10.1002/cne.902690207.

40. Jing, M., Li, Y., Zeng, J., Huang, P., Skirzewski, M., Kljakic, O., Peng, W., Qian, T., Tan, K., Zou, J., et al. (2020). An optimized acetylcholine sensor for monitoring in vivo cholinergic activity. Nat Methods 17, 1139–1146. 10.1038/s41592-020-0953-2.

41. Jones, I.W., Bolam, J.P., and Wonnacott, S. (2001). Presynaptic localisation of the nicotinic acetylcholine receptor beta2 subunit immunoreactivity in rat nigrostriatal dopaminergic neurones. J Comp Neurol 439, 235–247. 10.1002/cne.1345.

42. Kim, H.R., Malik, A.N., Mikhael, J.G., Bech, P., Tsutsui-Kimura, I., Sun, F., Zhang, Y., Li, Y., Watabe-Uchida, M., Gershman, S.J., and Uchida, N. (2020). A Unified Framework for Dopamine Signals across Timescales. Cell 183, 1600–1616 e1625. 10.1016/j.cell.2020.11.013.

43. Kimura, M., Rajkowski, J., and Evarts, E. (1984). Tonically discharging putamen neurons exhibit set- dependent responses. Proc Natl Acad Sci U S A 81, 4998–5001. 10.1073/pnas.81.15.4998.

44. Kosillo, P., Zhang, Y.F., Threlfell, S., and Cragg, S.J. (2016). Cortical Control of Striatal Dopamine Transmission via Striatal Cholinergic Interneurons. Cereb Cortex 26, 4160–4169. 10.1093/cercor/bhw252.

45. Kramer, P.F., Twedell, E.L., Shin, J.H., Zhang, R., and Khaliq, Z.M. (2020). Axonal mechanisms mediating gamma-aminobutyric acid receptor type A (GABA-A) inhibition of striatal dopamine release. Elife 9. 10.7554/eLife.55729.

46. Kudoh, S.N., and Taguchi, T. (2002). A simple exploratory algorithm for the accurate and fast detection of spontaneous synaptic events. Biosens Bioelectron 17, 773–782. 10.1016/s0956-5663(02)00053-2.

47. Lebowitz, J.J., Trinkle, M., Bunzow, J.R., Balcita-Pedicino, J.J., Hetelekides, S., Robinson, B., De La Torre, S., Aicher, S.A., Sesack, S.R., and Williams, J.T. (2021). Subcellular localization of D2 receptors in the murine substantia nigra. Brain Struct Funct. 10.1007/s00429-021-02432-3.

48. Lim, S.A., Kang, U.J., and McGehee, D.S. (2014). Striatal cholinergic interneuron regulation and circuit effects. Front Synaptic Neurosci 6, 22. 10.3389/fnsyn.2014.00022.

49. Liu, L., Zhao-Shea, R., McIntosh, J.M., Gardner, P.D., and Tapper, A.R. (2012). Nicotine persistently activates ventral tegmental area dopaminergic neurons via nicotinic acetylcholine receptors containing alpha4 and alpha6 subunits. Mol Pharmacol 81, 541–548. 10.1124/mol.111.076661.

50. Mamaligas, A.A., Cai, Y., and Ford, C.P. (2016). Nicotinic and opioid receptor regulation of striatal dopamine D2-receptor mediated transmission. Sci Rep 6, 37834. 10.1038/srep37834.

51. McIntosh, J.M., Azam, L., Staheli, S., Dowell, C., Lindstrom, J.M., Kuryatov, A., Garrett, J.E., Marks, M.J., and Whiteaker, P. (2004). Analogs of alpha-conotoxin MII are selective for alpha6-containing nicotinic acetylcholine receptors. Mol Pharmacol 65, 944–952. 10.1124/mol.65.4.944.

52. Mohebi, A., and Berke, J.D. (2020). Dopamine release drives motivation, independently from dopamine cell firing. Neuropsychopharmacology 45, 220. 10.1038/s41386-019-0492-7.

53. Mohebi, A., Pettibone, J.R., Hamid, A.A., Wong, J.T., Vinson, L.T., Patriarchi, T., Tian, L., Kennedy, R.T., and Berke, J.D. (2019). Dissociable dopamine dynamics for learning and motivation. Nature 570, 65–70. 10.1038/s41586-019-1235-y.

54. Obermayer, J., Luchicchi, A., Heistek, T.S., de Kloet, S.F., Terra, H., Bruinsma, B., Mnie-Filali, O., Kortleven, C., Galakhova, A.A., Khalil, A.J., et al. (2019). Prefrontal cortical ChAT-VIP interneurons provide local excitation by cholinergic synaptic transmission and control attention. Nat Commun 10, 5280. 10.1038/s41467-019-13244-9.

55. Olah, V.J., Tarcsay, G., and Brunner, J. (2021). Small Size of Recorded Neuronal Structures Confines the Accuracy in Direct Axonal Voltage Measurements. eNeuro 8. 10.1523/ENEURO.0059-21.2021.

56. Rama, S., Zbili, M., and Debanne, D. (2018). Signal propagation along the axon. Curr Opin Neurobiol 51, 37–44. 10.1016/j.conb.2018.02.017.

57. Ramon y Cajal, S. (1899). Textura Del Sistema Nervioso Del Hombre Y De Los Vertebrados: Estudios Sobre El Plan Estructural Y Composición Histológica De Los Centros Nerviosos Adicionados De Consideraciones Fisiológicas Fundadas En Los Nuevos Descubrimentos (Moya).

58. Raz, A., Feingold, A., Zelanskaya, V., Vaadia, E., and Bergman, H. (1996). Neuronal synchronization of tonically active neurons in the striatum of normal and parkinsonian primates. J Neurophysiol 76, 2083–2088. 10.1152/jn.1996.76.3.2083.

59. Rice, M.E., and Cragg, S.J. (2004). Nicotine amplifies reward-related dopamine signals in striatum. Nat Neurosci 7, 583–584. 10.1038/nn1244.

60. Ritzau-Jost, A., Tsintsadze, T., Krueger, M., Ader, J., Bechmann, I., Eilers, J., Barbour, B., Smith, S.M., and Hallermann, S. (2021). Large, Stable Spikes Exhibit Differential Broadening in Excitatory and Inhibitory Neocortical Boutons. Cell Rep 34, 108612. 10.1016/j.celrep.2020.108612.

61. Rothman, J.S., and Silver, R.A. (2018). NeuroMatic: An Integrated Open-Source Software Toolkit for Acquisition, Analysis and Simulation of Electrophysiological Data. Front Neuroinform 12, 14. 10.3389/fninf.2018.00014.

62. Salminen, O., Drapeau, J.A., McIntosh, J.M., Collins, A.C., Marks, M.J., and Grady, S.R. (2007). Pharmacology of alpha-conotoxin MII-sensitive subtypes of nicotinic acetylcholine receptors isolated by breeding of null mutant mice. Mol Pharmacol 71, 1563–1571. 10.1124/mol.106.031492.

63. Sethuramanujam, S., Matsumoto, A., deRosenroll, G., Murphy-Baum, B., Grosman, C., McIntosh, J.M., Jing, M., Li, Y., Berson, D., Yonehara, K., and Awatramani, G.B. (2021). Rapid multi-directed cholinergic transmission in the central nervous system. Nat Commun 12, 1374. 10.1038/s41467-021-21680-9.

64. Shin, J.H., Adrover, M.F., and Alvarez, V.A. (2017). Distinctive Modulation of Dopamine Release in the Nucleus Accumbens Shell Mediated by Dopamine and Acetylcholine Receptors. J Neurosci 37, 11166–11180. 10.1523/JNEUROSCI.0596-17.2017.

65. Shu, Y., Duque, A., Yu, Y., Haider, B., and McCormick, D.A. (2007). Properties of action-potential initiation in neocortical pyramidal cells: evidence from whole cell axon recordings. J Neurophysiol 97, 746–760. 10.1152/jn.00922.2006.

66. Sulzer, D., Cragg, S.J., and Rice, M.E. (2016). Striatal dopamine neurotransmission: regulation of release and uptake. Basal Ganglia 6, 123–148. 10.1016/j.baga.2016.02.001.

67. Sun, Y.G., Pita-Almenar, J.D., Wu, C.S., Renger, J.J., Uebele, V.N., Lu, H.C., and Beierlein, M. (2013). Biphasic cholinergic synaptic transmission controls action potential activity in thalamic reticular nucleus neurons. J Neurosci 33, 2048–2059. 10.1523/JNEUROSCI.3177-12.2013.

68. Susaki, E.A., Tainaka, K., Perrin, D., Yukinaga, H., Kuno, A., and Ueda, H.R. (2015). Advanced CUBIC protocols for whole-brain and whole-body clearing and imaging. Nat Protoc 10, 1709–1727. 10.1038/nprot.2015.085.

69. Szapiro, G., and Barbour, B. (2007). Multiple climbing fibers signal to molecular layer interneurons exclusively via glutamate spillover. Nat Neurosci 10, 735–742. 10.1038/nn1907.

70. Tanimura, A., Du, Y., Kondapalli, J., Wokosin, D.L., and Surmeier, D.J. (2019). Cholinergic Interneurons Amplify Thalamostriatal Excitation of Striatal Indirect Pathway Neurons in Parkinson’s Disease Models. Neuron 101, 444–458 e446. 10.1016/j.neuron.2018.12.004.

71. Tanimura, A., Pancani, T., Lim, S.A.O., Tubert, C., Melendez, A.E., Shen, W., and Surmeier, D.J. (2018). Striatal cholinergic interneurons and Parkinson’s disease. Eur J Neurosci 47, 1148–1158. 10.1111/ejn.13638.

72. Threlfell, S., Lalic, T., Platt, N.J., Jennings, K.A., Deisseroth, K., and Cragg, S.J. (2012). Striatal dopamine release is triggered by synchronized activity in cholinergic interneurons. Neuron 75, 58–64. 10.1016/j.neuron.2012.04.038.

73. Wang, L., Zhang, X., Xu, H., Zhou, L., Jiao, R., Liu, W., Zhu, F., Kang, X., Liu, B., Teng, S., et al. (2014). Temporal components of cholinergic terminal to dopaminergic terminal transmission in dorsal striatum slices of mice. J Physiol 592, 3559–3576. 10.1113/jphysiol.2014.271825.

74. Wonnacott, S. (1997). Presynaptic nicotinic ACh receptors. Trends Neurosci 20, 92–98. 10.1016/s0166-2236(96)10073-4.

75. Wu, J., George, A.A., Schroeder, K.M., Xu, L., Marxer-Miller, S., Lucero, L., and Lukas, R.J. (2004). Electrophysiological, pharmacological, and molecular evidence for alpha7-nicotinic acetylcholine receptors in rat midbrain dopamine neurons. J Pharmacol Exp Ther 311, 80–91. 10.1124/jpet.104.070417.

76. Yorgason, J.T., Zeppenfeld, D.M., and Williams, J.T. (2017). Cholinergic Interneurons Underlie Spontaneous Dopamine Release in Nucleus Accumbens. J Neurosci 37, 2086–2096. 10.1523/JNEUROSCI.3064-16.2017.

77. Zhang, H., and Sulzer, D. (2004). Frequency-dependent modulation of dopamine release by nicotine. Nat Neurosci 7, 581–582. 10.1038/nn1243.

78. Zhou, F.M., Liang, Y., and Dani, J.A. (2001). Endogenous nicotinic cholinergic activity regulates dopamine release in the striatum. Nat Neurosci 4, 1224–1229. 10.1038/nn769.

79. Zhou, F.M., Wilson, C.J., and Dani, J.A. (2002). Cholinergic interneuron characteristics and nicotinic properties in the striatum. J Neurobiol 53, 590–605. 10.1002/neu.10150.

80. Zoli, M., Moretti, M., Zanardi, A., McIntosh, J.M., Clementi, F., and Gotti, C. (2002). Identification of the nicotinic receptor subtypes expressed on dopaminergic terminals in the rat striatum. J Neurosci 22, 8785–8789.

